# The structural basis of lipid scrambling and inactivation in the endoplasmic reticulum scramblase TMEM16K

**DOI:** 10.1101/447417

**Authors:** Simon R. Bushell, Ashley C.W. Pike, Maria E. Falzone, Nils J. G. Rorsman, Chau M Ta, Robin A. Corey, Thomas D. Newport, Chitra A. Shintre, Annamaria Tessitore, Amy Chu, Qinrui Wang, Leela Shrestha, Shubhashish M.M. Mukhopadhyay, James D. Love, Nicola A. Burgess-Brown, Rebecca Sitsapesan, Phillip J. Stansfeld, Juha T. Huiskonen, Paolo Tammaro, Alessio Accardi, Elisabeth P. Carpenter

**Author notes:** These authors contributed equally to this work (Simon R. Bushell, Ashley C.W. Pike, Maria E. Falzone). These authors contributed equally to this work (Nils J. G. Rorsman, Chau M Ta, Robin A. Corey, Thomas D. Newport). These authors contributed equally to this work (Rebecca Sitsapesan, Phillip J. Stansfeld, Juha T. Huiskonen, Paolo Tammaro, Alessio Accardi). Nils Rorsman: OxSyBio, Atlas Building, Harwell Campus, Didcot, Oxfordshire, OX11 0QX, UK. Chau M. Ta, Department of Internal Medicine - Cardiovascular Medicine, University of Iowa, USA. Thomas D. Newport: Oxford Nanopore Technologies, Oxford Science Park, OX4 4DQ, UK. Chitra Shintre: Vertex Pharmaceuticals Ltd, Milton Park, Oxfordshire, UK. Annamaria Tessitore: Nuffield Division of Clinical Laboratory Sciences, Oxford University, UK. Amy Chu: Department of Biochemistry, Oxford University, UK. James Love: Institute for Protein Innovation, 4 Blackfan Circle, Boston, MA, USA.

## Abstract

Membranes in cells have defined distributions of lipids in each leaflet, controlled by lipid scramblases and flip/floppases. However, for some intracellular membranes such as the endoplasmic reticulum the scramblases have not been identified. Members of the TMEM16 family have either lipid scramblase and ion channel activity, or specific chloride channel activity. Although TMEM16K is widely distributed and associated with the neurological disorder autosomal recessive spinocerebellar ataxia type 10 (SCAR10), its location in cells, function and structure are largely uncharacterised. Here we show that TMEM16K is an ER-resident calcium-regulated lipid scramblase. Our crystal structures of TMEM16K show a scramblase fold, with an open lipid transporting groove. Additional structures solved by cryo-EM reveal extensive conformational changes extending from the cytoplasmic to the ER side of the membrane, giving a state with a closed lipid permeation pathway. Molecular dynamics simulations showed that the open-groove conformation is necessary for scramblase activity. Our results suggest mechanisms by which missense variants of TMEM16K could cause SCAR10 ataxia, providing new hypotheses to explore for therapy.

---

Cells and their organelles are enclosed by lipid bilayers and the lipid composition of either side of these membranes is controlled by active transporters (flippases and floppases) and passive scramblases, which equilibrate lipids between the membrane leaflets^1^. Many lipids are synthesized on the cytoplasmic side of the endoplasmic reticulum (ER) membrane which, unlike the plasma membrane (PM), has a symmetrical lipid distribution, suggesting a role for scramblases in the ER. To date, specific ER scramblases have not been identified and characterised. The ten members of the TMEM16 scramblase/channel family of integral membrane proteins show a surprising diversity of function, being either Ca^2+^-activated chloride channels (TMEM16A and B)^2–4^, or Ca^2+^-activated lipid scramblases with nonselective ion channel activity (TMEM16C, D, F, G and J)^5–8^. While some members of the family (A,B,F) reside in the plasma membrane, others, including TMEM16K^9^ may function in intracellular membranes. TMEM16K is a widely distributed^10^, but relatively unstudied member of the TMEM16K family. Truncations and missense variants of TMEM16K (ANO10) are associated with the autosomal recessive spinocerebellar ataxia SCAR10^11,12^ (as known as ARCA3^13–15^ or ATX-ANO10^16^). SCAR10 causing cerebellar ataxia, epilepsy and cognitive impairment with cerebellar atrophy noted on MRI brain and coenzyme Q10 deficiency found in muscle biopsy, fibroblasts and cerebrospinal fluid^11,12,17,18^. Some patients also have epilepsy and cognitive impairment^13,14^. Knockout studies in Drosophila^19^ and mice^20^ have suggested that loss of TMEM16K homologue function affects spindle formation^19^, Ca^2+^ signalling^20^ and apoptosis^19,20^.

Structural studies have gone some way towards explaining how TMEM16 family members function as channels or lipid scramblases. The crystal structure of the *Nectria haematococca* TMEM16 (nhTMEM16), a fungal lipid scramblase with non-selective channel activity, revealed a dimer arranged in a bi-lobal ‘butterfly’ fold, with each subunit containing a two Ca^2+^ ion binding site and ten transmembrane (TM) helices^21^. Each monomer has a hydrophilic, membrane-spanning groove that provides a route for lipid head groups to move across membranes. Molecular dynamics simulations subsequently confirmed this lipid scrambling mechanism *in silico*^22,23^. Structures of the mouse TMEM16A chloride channel revealed an alternative conformation, with two groove-associated transmembrane α helices blocking the top of the scramblase groove, forming a closed pore^24–26^.

Although patho-physiologically important, TMEM16K remains a poorly-characterized member of the TMEM16 family, as its cellular localization, function, regulation and structure are largely uncharacterised. Here we show that TMEM16K is an ER-resident calcium-activated lipid scramblase. We present structures of TMEM16K solved by both X-ray crystallography and cryo-electron microscopy, revealing a classic scramblase fold^21^, with extensive conformational changes propagated from the cytoplasmic to the ER face of the membrane, which lead to opening or closing of the lipid transporting groove. We use MD simulations to confirm that in TMEM16K the open groove conformation is necessary for scramblase activity. We also investigate the role of Ca^2+^-ion binding in conformational changes in TMEM16K.

## Results

### TMEM16K is an ER resident lipid scramblase

To test whether TMEM16K resides in intracellular membranes^27^ we explored its location in HEK293T cells using confocal microscopy. Under overexpression conditions, we saw nearly complete colocalisation between GFP-labelled TMEM16K and ER membranes stained with concanavalin A (Fig. 1a). Conversely, we saw little overlap in cells whose plasma membranes were stained with wheat germ agglutinin (Fig. 1b). We concluded that even under overexpression conditions TMEM16K is primarily confined to the ER in transfected cells. To test whether the small fraction of PM-resident TMEM16K functions as Ca^2+^-dependent ion channel we measured the whole-cell currents in HEK293T cells expressing TMEM16K in the presence of either 300 nM or 80 μM [Ca^2+^]_i_. These Ca^2+^ ion concentrations are sufficient to activate either TMEM16A, the PM chloride channel^2–4,6^, or TMEM16F, a lipid scramblase with non-selective ion channel activity^5,28^. Both these proteins are homologues of TMEM16K (Supplementary Fig. 1). We did not detect currents different from untransfected cells in either condition (Figs 1c,d), suggesting that TMEM16K does not mediate PM Ca^2+^-activated currents.

**Fig. 1.**
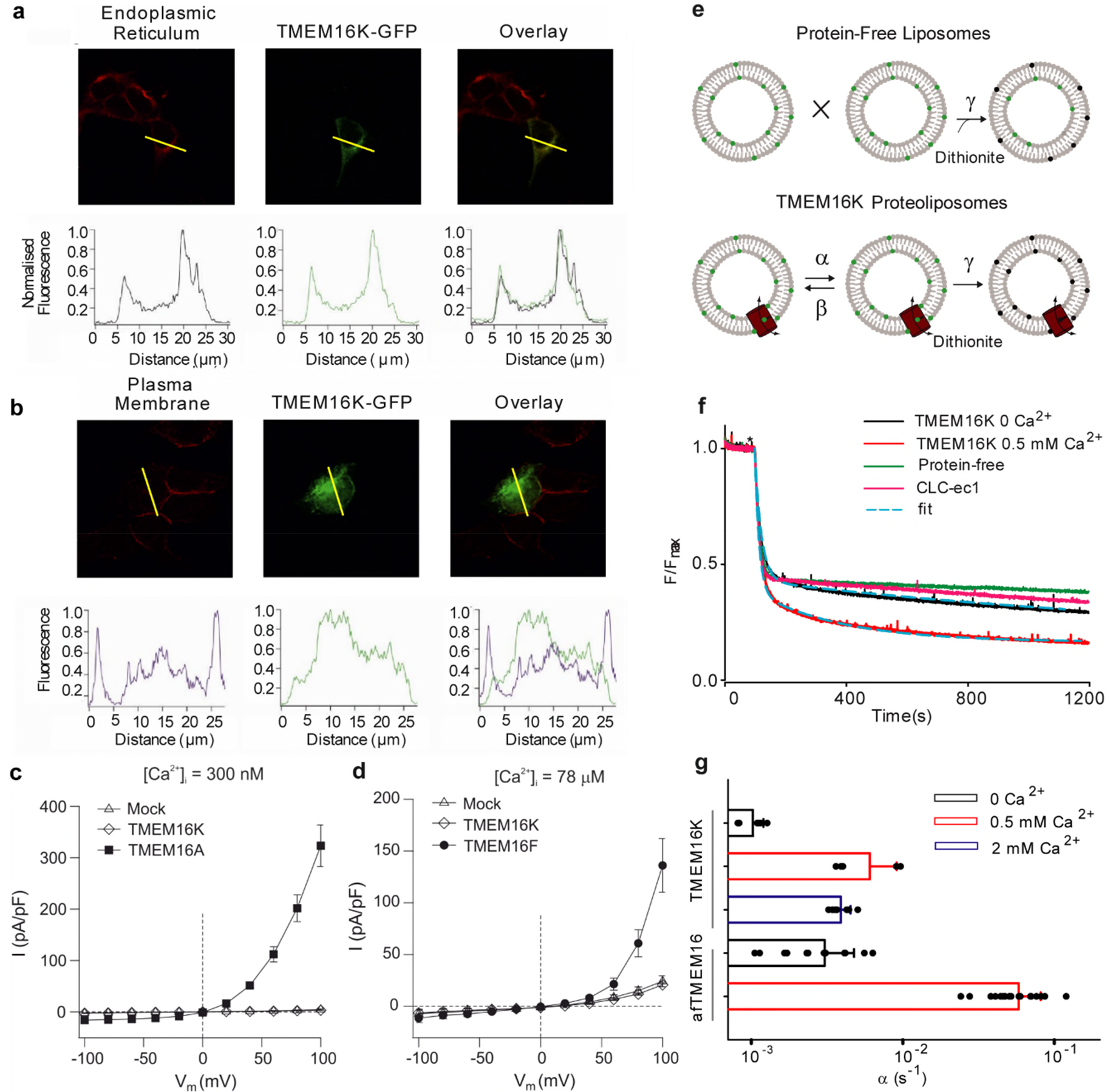
Characterisation of TMEM16K, an ER resident calcium activated scramblase. **a** Confocal images of HEK293T cells expressing TMEM16K-GFP (green). The ER was stained with AlexaFluor 594-concanavalin (red), as indicated. **b** Confocal images of HEK293T cells expressing TMEM16K-GFP (green). The PM was stained with AlexaFluor 555-Wheat Germ Agglutinin (red). The lower panels in **a** and **b** show the relative strength of the fluorescence signal along the yellow bar in the top panels. **c** Mean whole‐cell current *versus* voltage relationships for mock-transfected HEK‐293T cells (Mock, N=8) or cells expressing TMEM16A (N=8) or TMEM16K (N=6), as indicated. **d** Mean whole‐cell current *versus* voltage relationships for mock transfected HEK‐293T (Mock, N=5) cells or cells expressing TMEM16K (N=6) or TMEM16F (N=5), as indicated.**e** Schematic for the dithionite-based phospholipid scramblase activity. **f** Effect of Ca^2+^ on scramblase activity**. g** Bar chart showing the rate constants for the lipid scramblase activities for hTMEM16K and afTMEM16 with and without Ca^2+^.

Next, we used a dithionite-based scrambling assay^29^ (Fig. 1e,f) to investigate whether TMEM16K has Ca^2+^-regulated scramblase activity, as seen in mammalian TMEM16F^5,6,30^ and the fungal afTMEM16^29^ and nhTMEM16^21^ homologues. We found that purified hTMEM16K reconstituted into lipid vesicles transports lipids with a scrambling rate constant of ~0.003 s^−1^ in 0.5 or 2 mM Ca^2+^, corresponding to ~100 lipids translocated s^−1^ (Fig. 1f). Without Ca^2+^, the rate constant decreased ~4-fold, indicating that TMEM16K scramblase activity is Ca^2+^ regulated. Other TMEM16 scramblases, such as afTMEM16, are faster^30–32^, ~300-fold in 0.5 mM Ca^2+^ and ~15-fold without Ca^2+^ (Fig. 1g). Low scrambling rates in TMEM16K may be due to (i) a lack of residues equivalent to those involved in fast scrambling in the fungal homologues^8,22,32^; (ii) sensitivity to membrane composition; (iii) the absence of post-translational modifications or (iv) small molecules, proteins or lipid cofactors. Attempts to measure the ion channel activity with purified TMEM16K using either bulk flux assays with proteoliposomes^29,33^ or single channel planar lipid bilayer measurements^34^, were inconclusive, raising the question of whether this homologue also has the dual channel and scramblase activity, seen in other TMEM16 scramblases^5–7,28,29^.

We have shown that TMEM16K is an ER resident scramblase, and is therefore capable of providing the background scramblase activity required for maintaining the even distribution of lipids on either side of the ER membrane.

### Crystal structure determination and overall architecture

To understand the structural basis of lipid scrambling in TMEM16K we solved its structure by X-ray crystallography. We obtained initial diffraction data from vapour diffusion (VD) crystals of full-length human TMEM16K, which we improved by lipidic cubic phase crystallisation to obtain 3.4 Å resolution data. Both structures were solved by molecular replacement (Methods section, Supplementary Table 1). The structures were similar, although the crystal packing differs (Supplementary Fig. 2a,b,c), confirming that the conformation is not dictated by crystal contacts.

hTMEM16K is a symmetrical homodimer that adopts the classic TMEM16 butterfly fold ^21,24–26^ where each monomer comprises an N-terminal cytoplasmic domain (NCD) followed by a ten TM domain (TM1-TM10) and a C-terminal cytoplasmic α10 (Fig. 2a,b). The ER luminal surface of hTMEM16K is relatively compact, lacking the long, disulphide-bonded loops seen in mTMEM16A (Fig. 2a,b, Supplementary Fig. 2e). The N-terminal, cytoplasmic domain of hTMEM16K has a four-stranded β sheet (β1, β2, β3 and β8), with three helices on one side (α1, α3 and α4). These helices are covered by the β6-β7 hairpin insertion, which is unique to TMEM16K (Fig 2c,d). This β6-β7 hairpin and the associated loops provide important interactions between the cytoplasmic domain and the β9-β10 hairpin on the TM domain (Fig. 2d and Supplementary Fig. 3a). The other surface of the β-sheet is tightly packed against α8, part of an α-helical hairpin formed by α7 and α8, extensions of TM6 and TM7 (Supplementary Fig. 3b). This interaction links the position of the cytoplasmic domain to the TM helices. In the region between β3 and α3 the structures of hTMEM16K, mTMEM16A and nhTMEM16 differ significantly: hTMEM16K has a relatively short α3 and a β4-β5 hairpin that interacts directly with the C-terminal α10^A^ and α10^B^ helices (Supplementary Figs. 3c,d,e), linking the position of the cytoplasmic domain to the dimer interface helices.

**Fig. 2.**
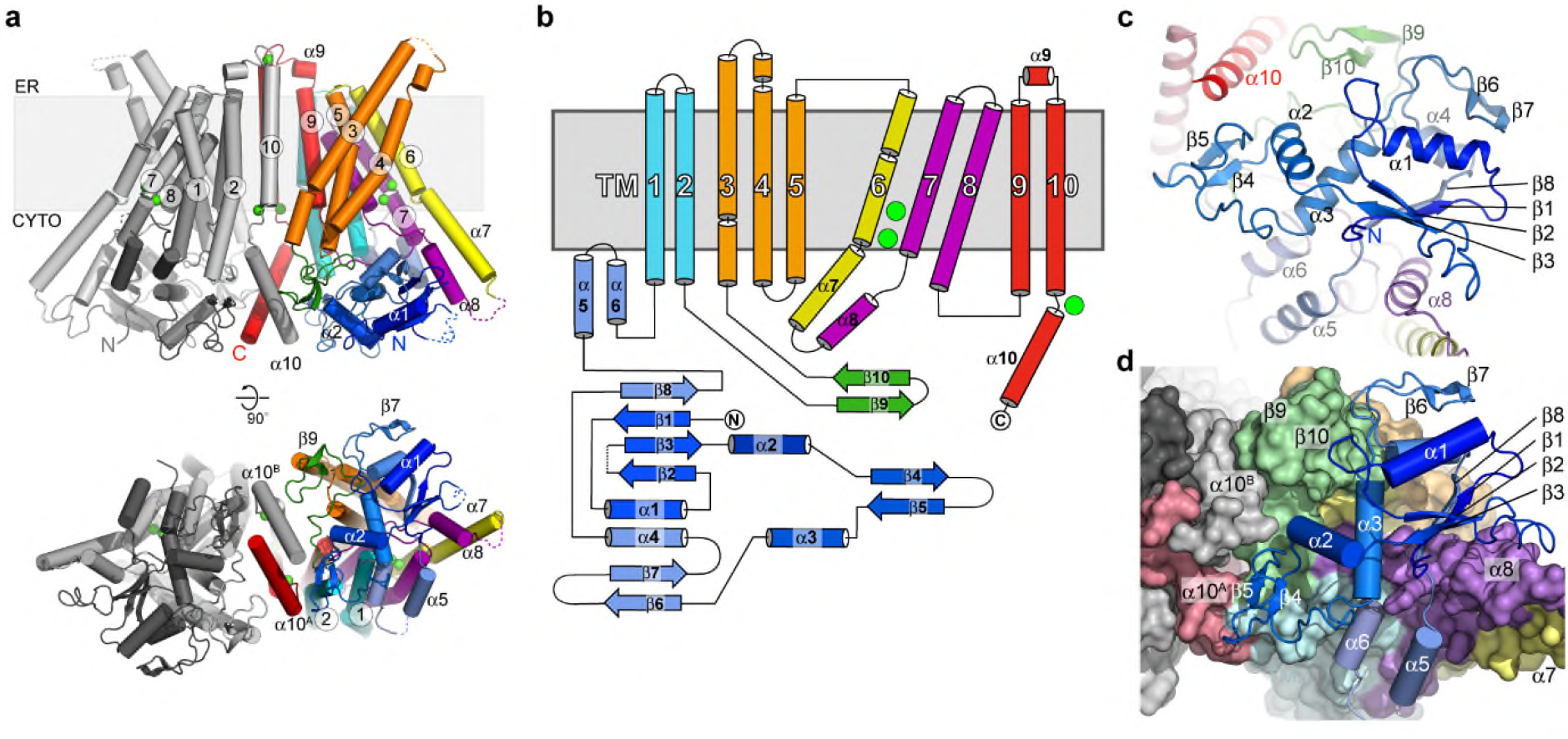
TMEM16K fold and cytoplasmic domain in the lipid scramblase conformation. **a** The structure of hTMEM16K viewed from the membrane plane and cytoplasm. In chain A the secondary structural elements are coloured as follows: the cytoplasmic domain (blue), TM1 and 2 (cyan), the β9-β10 hairpin (green), TM3, 4, 5 (orange), TM6 and α7 (yellow), α8, TM7 and 8 (purple), TM9, TM10 and α10 (red). Chain B is coloured grey. Ca^2+^ are coloured bright green. **b** TMEM16K secondary structure elements, coloured according to **a**,**c** Cytoplasmic domain (blue), α7 and α8 (yellow/purple) and α10 (red) of TMEM16K viewed from the cytoplasm.**d** Interactions of the cytoplasmic domain with the TM domain and the C-terminal helices (shown as a molecular surface coloured as in **a**), viewed from the cytoplasm.

The TM domain of TMEM16K is C-terminal to the cytoplasmic domain and has ten TM α helices with TM3 to TM7 forming the lipid scramblase “groove” and TM6 to TM8 forming the binding site for two Ca^2+^ ions, seen in other TMEM16s (Fig. 3a,b,c,d). In hTMEM16K, as in other TMEM16 scramblases, the groove is lined by a series of charged and hydrogen bonding sidechains that could interact with lipid headgroups (Fig. 3b).

**Fig. 3.**
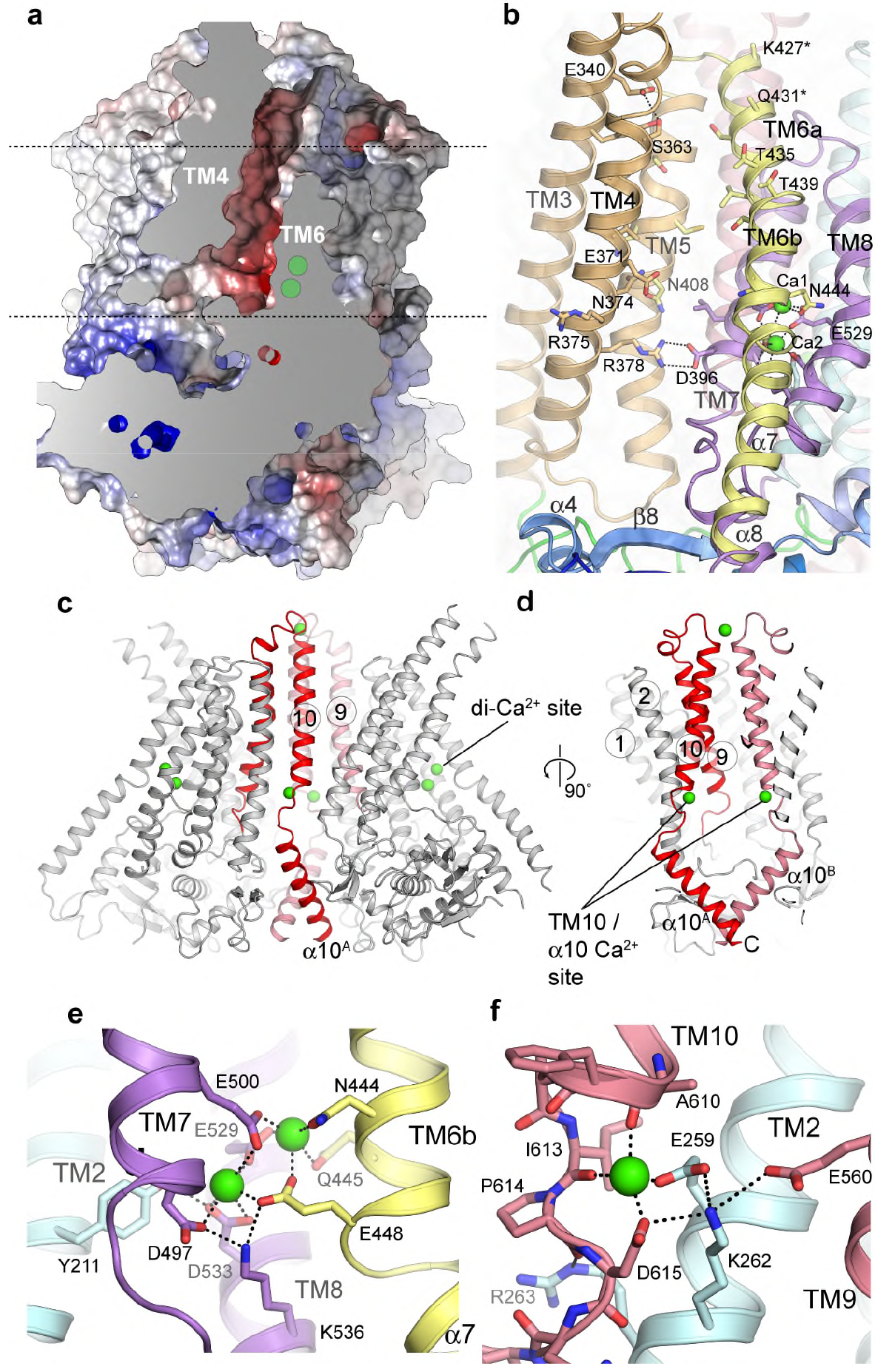
Structure of the TM domain region, the scramblase groove and the Ca^2+^ binding region. **a** Sliced molecular surface showing TMEM16K’s scramblase groove. Approximate positions of membrane and two Ca^2+^ ion site indicated by dotted lines and green circles. **b** Ribbon representation of the groove (coloured as per Figure 1b). Selected residues that were in contact with the lipid headgroups in MD simulations are shown as sticks and labelled.**c**, Location of the Ca^2+^-binding sites (green) in TMEM16K, and the dimer interface formed by TM10 and α10 contains a second Ca^2+^-binding site. **c,d** Ribbon representation of dimer highlighting positions of TM10-α10 helices (red). Zoomed in view of **e** the TM6-8 two Ca^2+^-binding site and of **f** the TM10/ α10 Ca^2+^-binding site.

We used molecular dynamics (MD) simulations to investigate how lipids traverse the TMEM16K groove. Coarse-grained (CG) MD simulations allowed us to sample the longer time scales necessary to investigate lipid scrambling events. nhTMEM16 is one of the few proteins to reveal lipid scrambling, within the >3500 membrane proteins in MemProtMD ^23^. CGMD simulations for hTMEM16K revealed lipid translocation events in both directions through the groove (Fig 4a,b,c,d). We also observed lipid scrambling upon conversion of intermediate states from the CG simulations to an atomistic description (Fig 4d), with lipids sampling the full translocation pathway. Taken together these results indicate that the TMEM16K groove serves as the lipid translocation pathway.

**Fig. 4.**
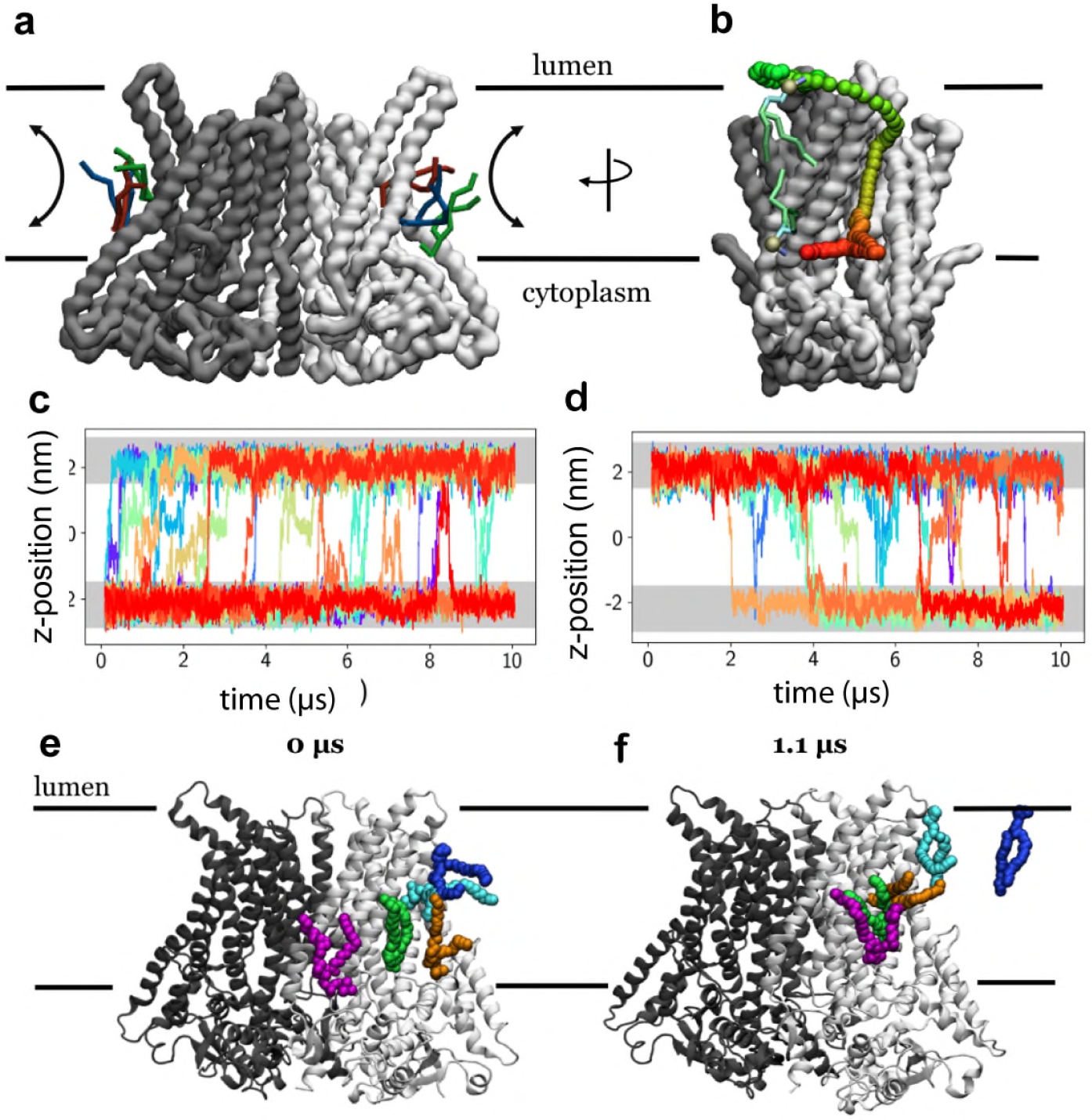
CG and all-atom molecular dynamics simulations of TMEM16K structures reveal lipid scrambling *in silico*. **a** View of CG TMEM16K after ~8 μs simulation. The protein backbone surface (gray and white) and 2-3 scrambling lipid molecules (blue, red and green sticks) are shown, **b** as panel a, but rotated by 90o and showing the pathway of a single lipid. The before and after poses are shown as green, blue and cyan sticks, with the head group particle shown as a gold sphere. The smoothed translocation head group pathway is denoted by red-to-green coloured spheres. **c,d** Traces showing quantification of CG lipid scrambling, across the z-axis, with the position of phosphate beads in each membrane leaflet shown in light grey. The traces show **c**, lipids that flip from cytoplasmic to lumenal leaflets, and **d** those that flip contrawise. **e,f** lipids scrambling through TMEM16K in atomistic simulation, coloured as per panel **a**.

The C-terminal region of TMEM16K forms the dimer interface, consisting of TM10 in the membrane region and its domain-swapped, cytoplasmic extension, α10 (Fig. 3c, Supplementary Fig. 4a-f). There is a large dimer interface cavity between TMs 3, 5, 9, 10 and TM10^B^, which contains several MAG7.9 lipids in the LCP structure (Supplementary Fig. 4e). In afTMEM16 we proposed that this cavity could play a role in deformation of the membrane^35^, it remains to be seen if it plays a similar role in TMEM16K.

### TMEM16K has both the TM6-8 two-Ca^2+^-binding site and an additional TM10-α10 Ca^2+^-binding site

Regulation of TMEM16 proteins by cytosolic Ca^2+^ is thought to involve the conserved two Ca^2+^ binding site situated between TM6, TM7 and TM8. This site is preserved in hTMEM16K (Fig. 3a-d), with anomalous difference maps confirming the presence of two Ca^2+^ ions (Supplementary Fig. 5a,b), and is formed from the sidechains of a series of highly conserved residues (Fig. 3e, Supplementary Fig. 1) and mutation of these residues reduced Ca^2+^ binding and stability in mTMEM16K^36^.

Surprisingly, our anomalous difference electron density maps revealed an additional Ca^2+^ ion bound near the dimer interface between TM2, TM10 and α10, where TM10 emerges from the membrane (Fig. 3c,d,f, Supplementary Fig. 5a,c). This Ca^2+^ ion interacts with the sidechains of Glu259 and Asp615 and the backbone carbonyls of Ala610 and Ile613, terminating the TM10 helix (Fig. 3e). In one case of ataxia associated with compound heterozygous TMEM16K mutations, one TMEM16K allele has an Asp615Asn mutation^12^, which is likely to impair Ca^2+^-binding at the TM10-α10 site (Fig. 3f), highlighting the importance of this site in TMEM16K function. Interestingly, inspection of the mTMEM16A electron density maps showed density at this location consistent with a Ca^2+^ ion (Supplementary Fig. 5d,e), and sequence conservation suggests this feature may be common to mammalian, but not fungal, TMEM16s (Supplementary Fig. 1).

### Cryo-EM structures of TMEM16K in low Ca^2+^ reveal a closed groove state

To understand whether TMEM16K can adopt alternative conformations and how changes in [Ca^2+^] affect the structure, we obtained electron cryomicroscopy (cryo-EM) structures for TMEM16K in 500 nM Ca^2+^ and in 10 mM EGTA (i.e. nominal Ca^2+^-free conditions), at resolutions of 4.5 Å and 5.3 Å, respectively (Supplementary Figs. 6 and 7, Supplementary Table 2). Surprisingly, in the presence of 500 nM Ca^2+^ the structure showed a dramatic change in conformation with extensive movements from the cytoplasmic to the ER side of the membrane, allowing a change from the classic scramblase open-groove arrangement described above, to a state where the groove is closed at the ER end. In the groove region, this conformation resembles the Ca^2+^-bound state of the mTMEM16A channel^24–26^ (Fig. 5a, Supplementary Fig. 8a). However, in the case of TMEM16K, these changes are accompanied by extensive changes that extend from the cytoplasmic to the ER side of the membrane, changes that were not observed in other TMEM16 structures. Although the Ca^2+^ concentration is reduced from 100 mM to 500 nM between the X-ray and this cryo-EM structures, there is clear electron density for Ca^2+^ ions at both the TM6-8 site and the TM10-α10 site (Supplementary Fig. 5d,e), indicating that these conformational changes are not induced by loss of Ca^2+^ binding at either site at the lower Ca^2+^ concentration.

**Fig. 5.**
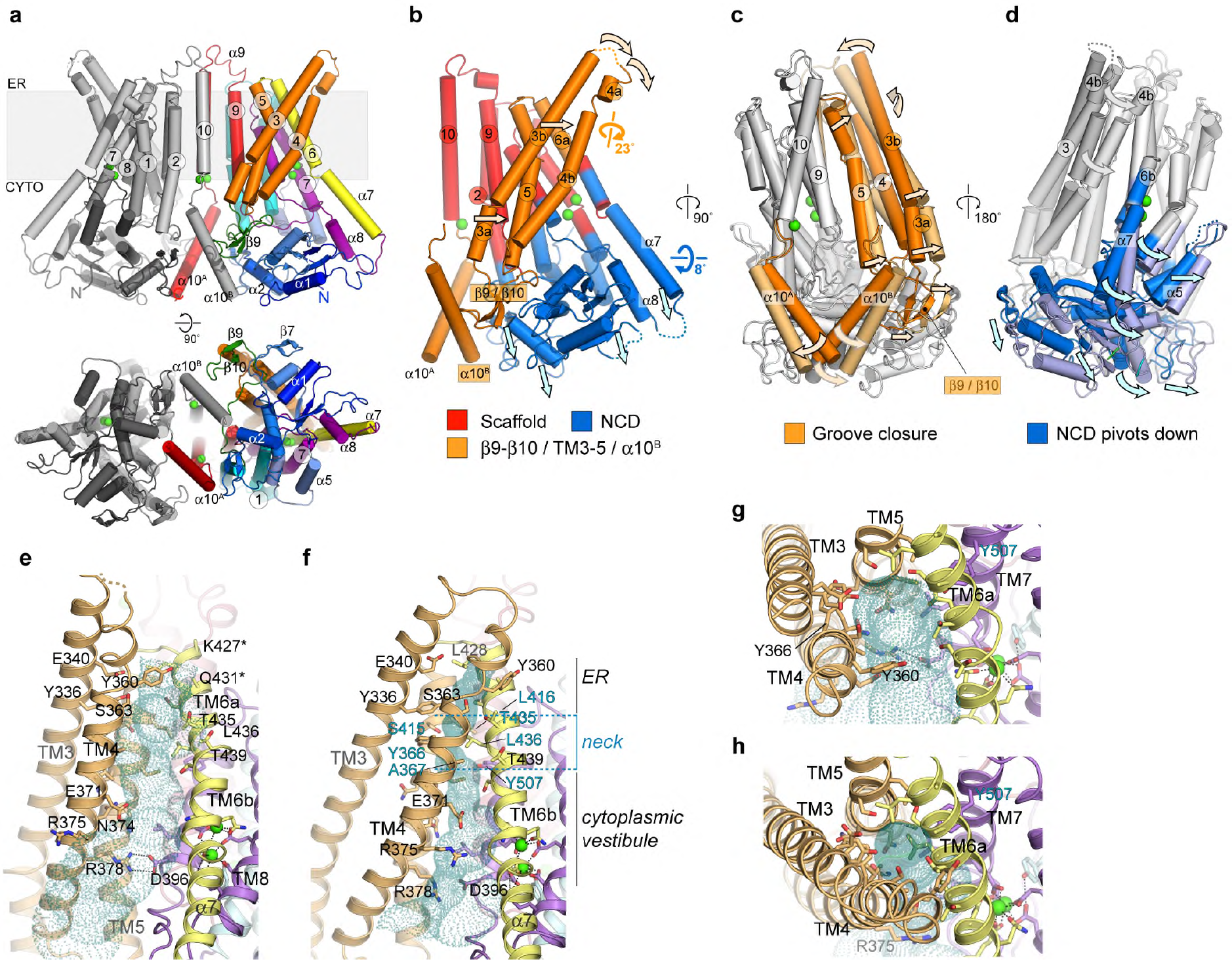
Cryo-EM structure of TMEM16K reveals structural changes from an open to a closed scramblase conformation. **a** Structure of TMEM16K in 500nM Ca^2+^ obtained by cryo-EM. Structural elements are coloured and viewed as in Fig. 2a. **b** Scaffold (red), TM4-6/α10B (orange) and NCD/α7-α8 regions (blue) mapped onto the monomer fold as derived from domain motion analysis. **c,d** Relative motions of **c**, the TM4-6/α10B region (orange) and **d**, the NCD/α7-α8 region, with the domains from the open groove crystal (dark colours) and closed groove 500nM Ca^2+^ cryo-EM (lighter colours) structures viewed looking onto the (**c**) dimer interface and (**d**) scramblase groove. **e-h**. Schematic representation of the scramblase groove in the crystal structure (**e**, **g**) and 500nM Ca^2+^ structure (**f**, **h**). The groove is viewed perpendicular to the membrane normal (**e**, **f**) and from the ER luminal face (**g, h**). Pale green dotted surface represents the groove/channel profile as calculated by HOLE^66^.

The change from an open to a closed scramblase groove state is associated with extensive movements of the cytoplasmic domain and the TM3-5 region, leading to the ER end of the TM3-TM4 α-helical hairpin blocking the ER end of the groove (Fig. 5, Supplementary Fig. 8). Within the TMEM16K monomer two large-scale movements occur between the open and closed groove structures, shown by torsional analysis using DYNDOM^37^ (Fig. 5b,c,d, Methods section and Supplementary Fig. 8, Supplementary Video). There is a rigid scaffold structure consisting of TM1-2, TM7-8 and TM9-10, the central section of TM6 (TM6b) and the two Ca^2+^ ions (Fig. 5b). The rest of the structure is divided into two parts that move separately. The first unit, consisting of the N-terminal cytoplasmic domain up to α6, and the α7-α8 helices (cytoplasmic extensions of TM6 and 7), rotates 8° relative to the scaffold structure (Fig. 5d). This causes the cytoplasmic domain to move down and away from the TM helices. The second unit, referred to here as the TM3-5 unit, consists of the β9-β10 hairpin, TM3, TM4 and TM5, as well as α10^B^. This whole unit rotates by ~23° relative to the scaffold domain, bringing α10 and the β9-β10 hairpin toward the front of cytoplasmic domain. This allows TM3, TM4 and TM5 to move up and pivot, so that TM4 fills the top of the groove (Fig. 5c,f,h). TM6 lies between the scaffold and the TM3-5 unit, and is divided into TM6a, which moves with the TM3-5 unit, and TM6b, which forms part of the two Ca^2+^-binding site and is associated with the scaffold domain (Supplementary Fig. 8g). TM6 therefore provides a flexible buffer between the scaffold and the TM3-5 unit, and has an ordered α7 extension which interacts with α8 and the cytoplasmic domain (Supplementary Fig. 8c). At the dimer interface, although the TM10 contacts are similar, the α10^A^-α10^B^ contact in the cytoplasm is lost completely in the cryo-EM structure, so the overall dimer interface area decreases from 1760 Å^2^ to 920 Å^2^ (Supplementary Fig. 4f-i). Interestingly, the TM3-5 unit forms one side of the hydrophobic dimer cavity which could provide space to accommodate these movements, potentially even allowing greater opening of the groove, as recently proposed^38^, to accommodate PEGylated lipids^39^.

In the nhTMEM16 open scramblase structure, the β9-β10 hairpin and α10^B^ positions resemble those in the open TMEM16K: both are pulled down beneath the TM domain, pivoting the TM3-5 unit to open the scramblase groove. In contrast, in the mTMEM16A channel the β9-β10 hairpin is replaced by a large disordered insertion (Supplementary Fig. 1), so TMEM16A may be unable to undergo the full pivoting movement of TM3-5 that is necessary to open the groove into the scramblase conformation. In addition to these large-scale changes, there are smaller movements, such as changes in the conformation of the ER ends of TM3 and TM4 allowing TM4 to wrap over the groove (Supplementary Fig. 8g-h).

The closed scramblase groove has three distinct sections, a hydrophilic vestibule on the ER side of the membrane, a narrow neck and a cytoplasmic vestibule (Fig 5f,h). The hydrophobic neck is lined with residues from TM4-7 (Tyr366, Ala367, Leu416, Ser415, Thr435, Leu436 and Tyr507), which together form a physical barrier to lipid movement (Fig. 5f,h). Unsurprisingly, we did not observe lipid scrambling in CGMD simulations of TMEM16K in this closed-groove conformation (Supplementary Fig. 8i).

### TMEM16K conformation in the absence of Ca^2+^

To investigate how Ca^2+^ binding regulates the TMEM16K conformation we determined the cryo-EM structure of TMEM16K in 10 mM EGTA to 5.3 Å resolution. This structure showed that the groove remains closed at the ER end as seen in the 500 nM Ca^2+^ structure and revealed additional rearrangements around both Ca^2+^-binding sites. The TM6-8 two Ca^2+^-binding site expands, with TM6b and TM8 moving away from TM7, and the C-terminal end of the α7 TM6 extension becoming disordered (Fig. 6a-d). These movements recapitulate the changes seen in afTMEM16^35^. The TM10-α10 Ca^2+^ binding site also changes, with TM9, α9 and TM10 moving down towards the cytoplasmic domain, and the TM1-loop-TM2 region moving away from TM10 and the dimer axis (Fig. 6d). The two Ca^2+^-binding site is integral to the scaffold domain and forms one side of the groove, so it is not surprising that conformational changes associated with loss of Ca^2+^ affect scramblase activity. However, TMEM16K retains basal scramblase activity even in the absence of Ca^2+^, suggesting that there is an additional, as yet unknown, open-groove Ca^2+^-free scramblase conformation (Fig 7).

**Fig. 6.**
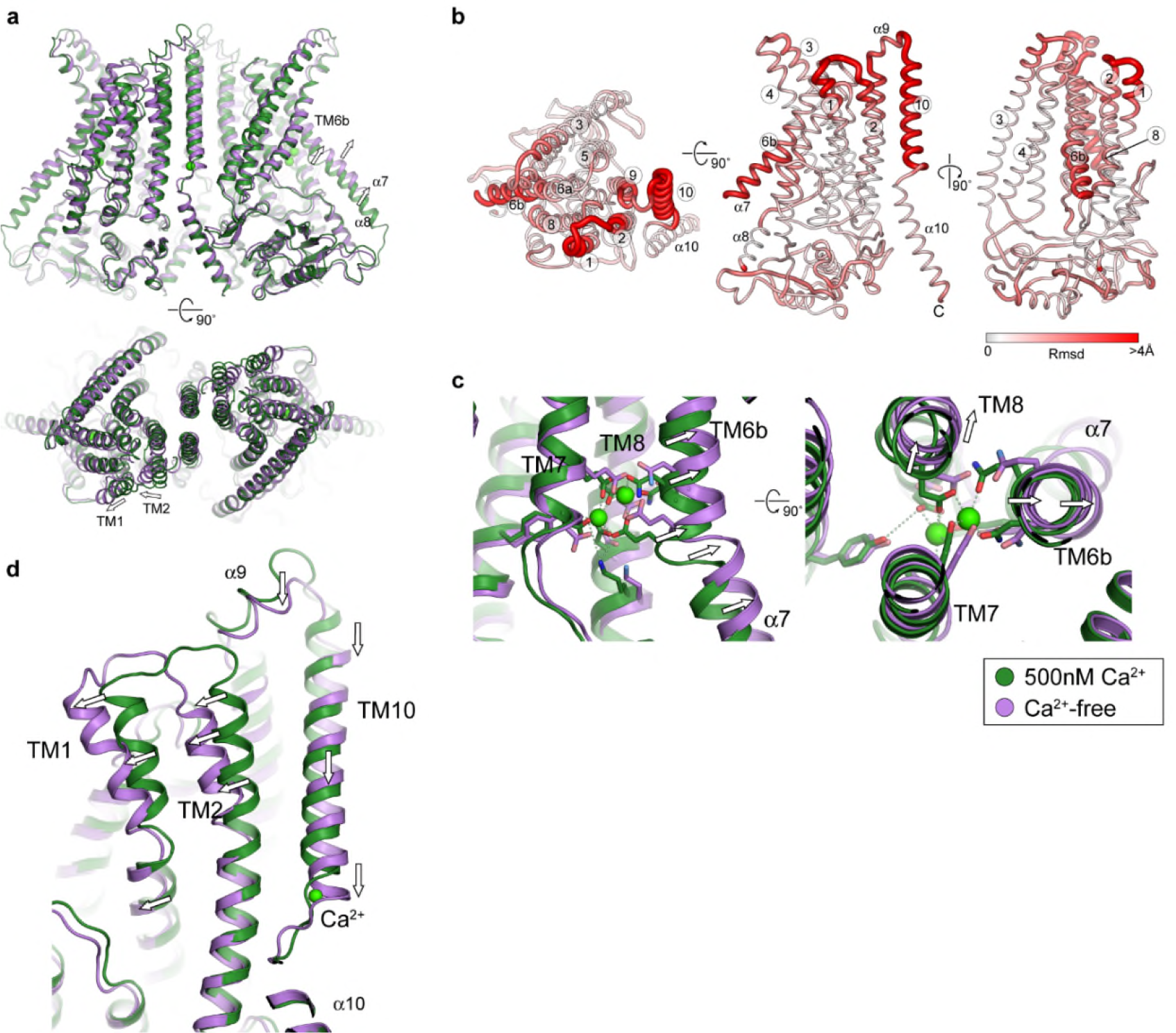
Comparison of the Ca^2+^-free and 500 nM cryo-EM structures of TMEM16K. **a** Overall structure of Ca^2+^-free state (purple), superimposed on the 500 nM Ca^2+^ structure (green). **b** Global conformational differences between 500nM and Ca^2+^-free structures. Schematic representations of the rms deviation (rmsd) in mainchain atomic positions mapped onto the monomer structure. Monomers from each structure were superposed using all atoms with LSQKAB (CCP4). The monomer is viewed from the luminal face (*left*), perpendicular to the membrane normal (*middle*) and scramblase groove face (*right*). The thickness and colour of the tube reflects the magnitude of the rmsd between the two structures. The main differences are localised in TM1/2, TM6b, α7, TM9 and TM10. **c** View of the two Ca^2+^-binding site, showing overall movement of TM6b / TM8 helices. **d** View of the TM10-α10 Ca^2+^-binding site, showing movement of TM1-2 and TM10 helices. TM9, α9 and TM10 move towards the cytoplasmic face of the membrane but α10 remains relatively fixed.

## Discussion

Our structures of TMEM16K have allowed us to explore the extensive conformational changes throughout the structure that occur when the lipid scramblase groove opens and closes (Fig 7). These changes are propagated from cytoplasmic to the ER face of the membrane through movements of two rigid bodies, one involving the α10^B^, β9-β10 hairpin and the TM3-TM5 unit and the other involving the cytoplasmic domain and the α7-α8 helices. The first unit rotates and moves up so that TM3-TM4 blocks the groove, while the second unit moves down to allow the first unit space to rotate. Surprisingly, these changes in TMEM16K conformation are not caused on changes in Ca^2+^ binding, as all three Ca^2+^-binding sites are occupied in both structures. TMEM16K scrambling is reduced in the absence of Ca^2+^ and complete removal of Ca^2+^ leads to further changes in the vicinity of the Ca^2+^ binding sites and at the lipid translocation pathway, explaining the reduction in activity.

**Fig. 7.**
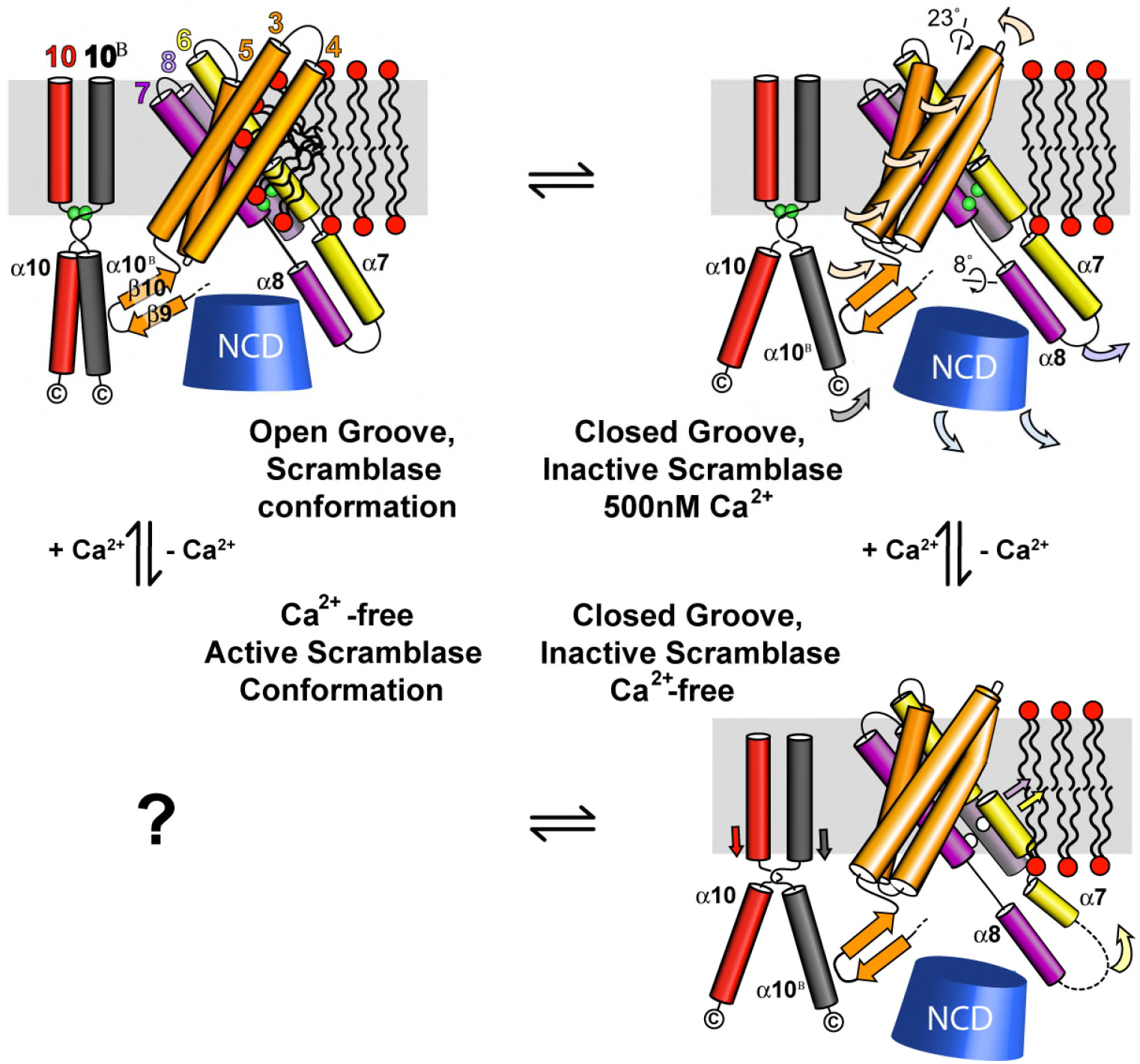
Schematic of conformational changes in TMEM16K. Schematic summary of TMEM16K conformational states, with the N-terminal cytoplasmic domain (NCD) shown as a blue cone and lipids with headgroups (red) and aliphatic chains (black).

We have demonstrated that TMEM16K resides in the ER, and has Ca^2+^-regulated scramblase activity, with a scrambling rate of ~100 lipids s^−1^. This relatively slow rate, with a modest 4-fold activation by Ca^2+^, could contribute to the background scrambling activity required to maintain the symmetrical lipid distribution in the ER membrane. Continuous scrambling and/or flip/floppase activities are needed in ER membranes to redistribute lipids synthesised on its cytoplasmic face. TMEM16K and other scramblases such as TMEM16G^27^ and the opportunistic GPCR scramblases^40–42^, could contribute to maintenance of this symmetry. In contrast, the PM’s highly asymmetrical lipid distribution is broken down only under extreme conditions such as apoptosis so that PM scramblases, like TMEM16F, may require tight regulation and high scrambling rates^5,6,30^. The structures of the human TMEM16K presented here, together with the conformational changes seen in afTMEM16^35^, demonstrate that there is extensive flexibility in TMEM16 structures, allowing them to adopt a range of activation states, potentially triggered by diverse stimuli such as lipid composition, post translational modification or cofactor binding.

TMEM16K has been associated with several cellular phenomena, including spindle formation^19^, Ca^2+^ signalling^20^, volume regulation^9^ and apoptosis^19,20^. Our work establishes its role as an ER scramblase, suggesting that the relationship between these cellular functions and lipid distributions warrants investigation.

These results provide a structural framework to understand how compound heterozygous missense mutations found in patients with SCAR10 ataxia could affect TMEM16K function. One mutation, Asp615Asn^12^, lies in the novel TM10-α10 Ca^2+^-binding site (Fig. 3g), where it forms one of the Ca^2+^ ion ligands and an ionic bond with Lys262, linking TM2 to the connection between TM10 and α10. There are also four missense mutations which introduce residues with very different sized side chains into key positions between α helices, likely disrupting helix packing (Gly229Trp^13^, Leu510Arg^11^)(Fig. 8d,c) or interfering with conformational changes (Phe171Ser^13^, Phe337Val^13^) (Fig. 8d,e). The observation that TMEM16K truncations and missense variants lead to SCAR10 and our observation that TMEM16K is an ER scramblase, suggest that the underlying cause of this ataxia could be associated with incorrect lipid distributions in ER and other membranes. This hypothesis provides a new direction for research into the underlying biology of these ataxias and the development of novel therapies.

**Fig. 8.**
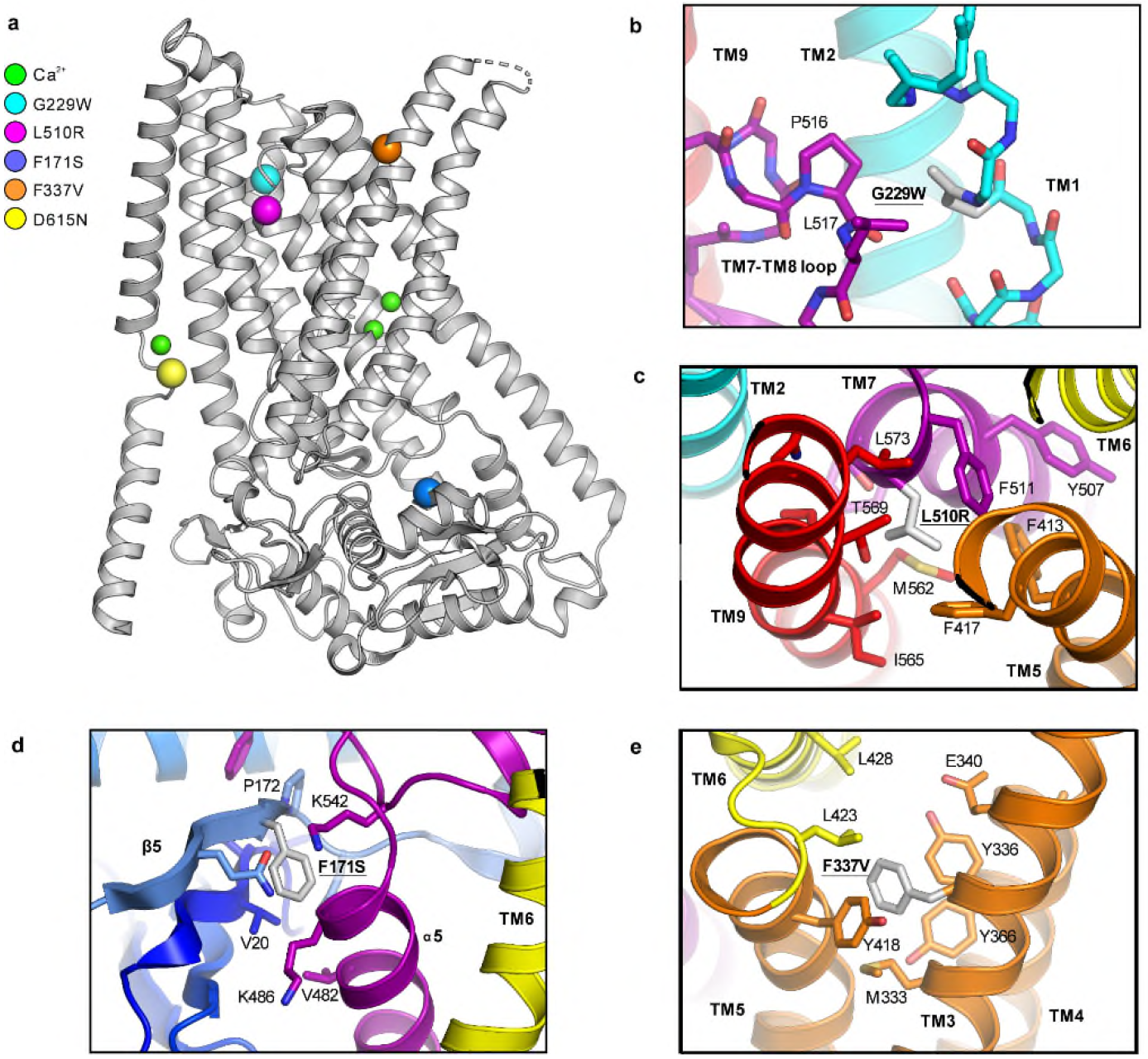
Missense variants in TMEM16K found in patients with SCAR10 ataxia. **a** Overview of TMEM16K with disease associated missense mutation locations shown as coloured spheres and Ca ^2+^ ions shown as smaller green spheres. Local environment for the disease associated mutations: **b** Gly229Trp, **c** Leu510Arg, **d** Phe171Ser and **e** Phe337Val.

## Methods

### Cell culture and transfection for chloride channel electrophysiology

TMEM16A (Genbank NM_178642), TMEM16F (NM_175344) and TMEM16K (NM_018075) subcloned into the pcDNA3.1 vector were used in this study. HEK-293T cells were cultured as previously described^43^ and transfected with 0.6μg of TMEM16A, TMEM16F or TMEM16K and 0.2μg of CD8 construct using Fugene HD (Promega, UK) according to the manufacturer’s instructions. Cells were used ~12-24 hours after transfection. Transfected cells were visualized using the anti-CD8 antibody-coated beads method^43^. Mock transfected cells refer to cells treated with transfection reagent only.

### Electrophysiology

TMEM16s currents were measured with the whole-cell configuration of the patch-clamp technique using an Axon 200B amplifier (Molecular Devices, USA) controlled with GE-pulse software (http://users.ge.ibf.cnr.it/pusch/programs-mik.htm). Currents were low-pass filtered at 2 kHz and sampled at 10 kHz. Pipettes were prepared from borosilicate glass capillary tubes (Harvard Apparatus, USA) using a PC-10 pipette puller (Narishige, Japan). Pipette tip diameter yielded a resistance of ~2-3 MΩ in the working solutions. The bath was grounded through a 3 M KCl agar bridge connected to a Ag-AgCl reference electrode. Experiments were conducted at 20-22 °C. The cell capacitance was assessed by measuring the area beneath a capacitive transient elicited by a 10 mV step or via the cell capacity compensation circuit of the amplifier. Current density was obtained by dividing the current amplitude for the cell capacitance.

The extracellular solution contained: 150 mM NaCl, 1 mM CaCl_2_, 1 mM MgCl_2_, 10 mM glucose, 10 mM D-mannitol and 10 mM HEPES; pH was adjusted to 7.4 with NaOH. The intracellular solution contained: 130 mM CsCl, 10 mM EGTA, 1 mM MgCl_2_, 10 mM HEPES and 8 mM CaCl_2_ to obtain [Ca^2+^]_i_ of ~0.3 μM; pH was adjusted to 7.3 with NaOH. The intracellular solution containing ~78 μM [Ca^2+^]_i_ was obtained by replacing EGTA with equimolar H-EDTA and by adding a total of 9 mM CaCl_2_.

### Membrane and ER staining

Staining of membrane compartments was performed on HEK-293T cells plated on 25 × 25 mm glass cover slips and transfected with TMEM16K tagged with GFP (TMEM16K-GFP). The cells were washed three times with cold PBS (Gibco, UK) and fixed in 4% paraformaldehyde (PFA) in PBS for 20 min. After fixation, the cells were washed three times with cold PBS and incubated with 5 μg/ml Wheat Germ Agglutinin, Alexa Fluor 555 Conjugate (ThermoFisher, UK) for 10 min in the dark at room temperature. The cover slips were then washed twice with cold PBS. For staining of the endoplasmic reticulum (ER), the cells were permeabilised with 0.2% (v/v) Triton-X-100 (Merck, UK) in PBS for 1 minute. Following this, the cells were washed with cold PBS and then incubated with 100 μg/ml concanavalin Alexa Fluor 594 Conjugate (Thermo Fisher, UK) in the dark for 30 min. The coverslips were then washed three times with PBS before being mounted onto microscope slides using a drop of Vectashield mounting medium (Vector Laboratories, USA) and sealed with generic nail varnish. The microscope slides were stored in the dark at 4 °C until imaged.

Cells were imaged using a LSM510 META confocal laser scanning system (Zeiss, UK) connected to an inverted AxioVert 200 microscope with a 63X objective (Zeiss, UK), controlled using the Zen 2012 (Blue edition) software (Zeiss, UK). TMEM16K-GFP expression and localisation was visualised using an excitation wavelength of 488 nm whilst emission at 509 nm was recorded. In the same samples, the ER or PM was visualised by excitation and emission wavelengths of 633 nm and 594 nm, or 544 nm and 555 nm, respectively. In all cases the fluorescence intensity of each cross section was normalised for the maximal intensity measured within the same cross section. The cellular distribution of TMEM16K-GFP in relation to the ER and PM stains was estimated via fluorescence intensity plots of cross sections taken from these images using the FIJI plug in package of Image J (U.S. National Institutes of Health).

### Cloning and expression of TMEM16K for structural and functional studies

The *Homo sapiens* TMEM16K gene, which encodes the TMEM16K/anoctamin-10 protein, was provided by the DNASU Plasmid collection (https://dnasu.org/DNASU/Home.do). The construct used for structure determination included the full length TMEM16K sequence, Met1 to Thr660, with a C-terminal purification tag with a tobacco etch virus (TEV) protease cleavage site, a 10x His purification sequence and a FLAG tag, in the expression vector pFB-CT10HF-LIC (available from The Addgene Nonprofit Plasmid Repository). Baculoviruses were produced by transformation of DH10Bac cells. *Spodoptera frugiperda* (Sf9) insect cells in Sf900II^TM^ serum free, protein free insect cell medium with L-glutamine (Thermofisher Scientific) were infected with recombinant baculovirus for virus amplification for 65 h at 27°C in 1 L shaker flasks. Baculovirus were harvested via centrifugation at 900 × *g* for 20 mins and the virus containing supernatant was used to infect 1 L of mid-log phase (2×10^6^ cells ml^−1^) Sf9 cells in Sf-900™ II Serum Free Medium (Gibco/Thermo-Fisher) in a 3 L flask, which were then grown for 72 h at 27 °C. Cells were harvested by centrifugation at 900 × *g* for 15 mins, washed with Phosphate Buffered Saline (PBS), and pelleted again prior to flash freezing in liquid N_2_, then stored at −80 °C until needed.

### TMEM16K purification

Cell pellets were resuspended in 30 ml / L equiv. original cell culture Buffer A (20mM HEPES pH 7.5, 200 mM NaCl, 5 % glycerol v/v, 2 mM tris(2-carboxyethyl)phosphine (TCEP) and lysed by two passes through a Emulsiflex C5 homogeniser (Avestin). n-Undecyl-β-D-Maltopyranoside (UDM) was added to a final concentration of 1 % (w/v) to the cell lysate, which was then incubated at 4 °C for 1 hr on a roller. Insoluble material was removed via centrifugation at 32,000 × *g* for 1 hr at 4 °C. The supernatant was supplemented with 5 mM final imidazole pH 7.5 before adding 50 % slurry (v/v) pre-equilibrated Talon^TM^ resin (1 ml slurry per L original culture volume). The suspension was then incubated at 4°C on a roller for 1 hr. Talon resin was collected by centrifugation at 900 × *g* for 15 mins, transferred to a gravity column where the remaining liquid was allowed to flow through. The resin was washed with 100 ml Buffer A supplemented with 20 mM imidazole, pH 7.5. Subsequently, all buffer solutions were supplemented with 0.045 % (w/v) UDM. TMEM16K was eluted with Buffer A supplemented with 250 mM imidazole. The protein’s C-terminal 10xHistidine-FLAG tag was removed by overnight incubation with 5:1 (w/w) TMEM16K : Tobacco Edge Virus (TEV) protease. TEV protease and non-cleaved contaminants were removed via reverse IMAC with Talon resin. Cleaved TMEM16K was concentrated to ~1ml and further purified by size exclusion chromatography on a Superose 6 Size exclusion Column in buffer A. When appropriate, free calcium concentration was determined using Calcium Green™-1 fluorescent indicator (Thermo-Fisher Scientific) according to the manufacturer’s instructions. Calcium concentration was calculated from a standard curve derived from solutions of known calcium concentration prepared using the Calcium Calibration Buffer Kit #1 (Thermo-Fisher Scientific) according to the manufacturer’s instructions.

### afTMEM16 Expression and Purification

afTMEM16 was expressed and purified as described^29^. Briefly, *S. cerevisiae* carrying pDDGFP2^44^ with afTMEM16 were grown in yeast synthetic drop-out medium supplemented with Uracil (CSM-URA; MP Biomedicals) and expression was induced with 2% (w/v) galactose at 30° for 22 hours. Cells were collected, snap frozen in liquid nitrogen, lysed by cryomilling (Retsch model MM400) in liquid nitrogen (3 × 3 min, 25 Hz), and resuspended in buffer A (150 mM KCl, 10% (w/v) glycerol, 50 mM Tris-HCl, pH8) supplemented with 1 mM EDTA, 5 μg ml^−1^ leupeptin, 2 μg ml^−1^ pepstatin, 100 μM phenylmethane sulphonylfluoride and protease inhibitor cocktail tablets (Roche). Protein was extracted using 1% (w/v) digitonin (EMD biosciences) at 4°C for 2 hours and the lysate was cleared by centrifugation at 40,000 g for 45 minutes. The supernatant was supplemented with 1 mM MgCl_2_ and 10 mM Imidazole, loaded onto a column of Ni-NTA agarose resin (Qiagen), washed with buffer A + 30 mM Imidazole and 0.12% digitonin, and eluted with buffer A + 300 mM Imidazole and 0.12% digitonin. The elution was treated with Tobacco Etch Virus protease overnight to remove the His tag and then further purified on a Superdex 200 10/300 GL column equilibrated with buffer A supplemented with 0.12% digitonin (GE Lifesciences). The afTMEM16 protein peak was collected and concentrated using a 50 kDa molecular weight cut off concentrator (Amicon Ultra, Millipore).

### Liposome reconstitution and lipid scrambling assay

Liposomes were prepared as described^29^ from a 7:3 mixture of 1-palmitoyl-2-oleoyl-glycero-3-phosphocholine (POPC) 1-palmitoyl-2-oleoyl-glycero-3-phosphoglycerol (POPG). Lipids in chloroform (Avanti), including 0.4% w/w tail labelled NBD-PE, were dried under N_2_, washed with pentane and resuspended at 20 mg ml^−1^ in buffer B (150 mM KCl, 50 mM HEPES pH 7.4) with 35 mM 3-[(3-cholamidopropyl)dimethylammonio]-1-propanesulfonate (CHAPS). TMEM16K or afTMEM16 was added at 5 μg protein/mg lipids and detergent was removed using five changes of 150 mg mL^−1^ Bio-Beads SM-2 (Bio-Rad) with rotation at 4°C. Calcium or EGTA were introduced using sonicate, freeze, and thaw cycles. Liposomes were extruded through a 400 nm membrane and 20 μL were added to a final volume of 2 mL of buffer B supplemented with 0.5 or 2 mM CaCl_2_ or 2 mM EGTA. The fluorescence intensity of the NBD (excitation-470 nm emission-530 nm) was monitored over time with mixing in a PTI spectrophotometer and after 100s sodium dithionite was added at a final concentration of 40 mM. Data was collected using the FelixGX 4.1.0 software at a sampling rate of 3 Hz.

### Quantification of scrambling activity

Quantification of the scrambling rate constants of TMEM16K and afTMEM16 were determined as recently described ^33^, with the additional assumption that the forward and reverse rate constants for scrambling are the same. Briefly, the fluorescence time course was fit to the following equation

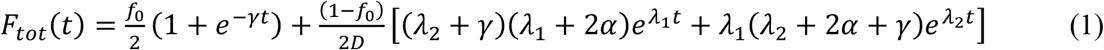

Where F_tot_(t) is the total fluorescence at time t, LiPF is the fraction of NBD-labelled lipids in the inner leaflet of protein free liposomes, γ=γ’[D] where γ’ is the second order rate constant of dithionite reduction, [D] is the dithionite concentration, f_0_ is the fraction of protein-free liposomes in the sample, α is the scrambling rate constant and

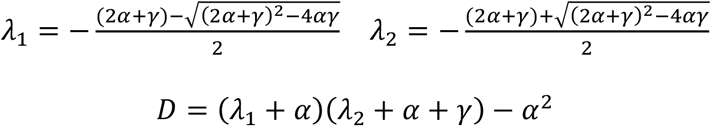

The free parameters of the fit are f_0_ and α while LiPF and γ are experimentally determined from experiments on protein-free liposomes. In protein-free vesicles a very slow fluorescence decay is visible likely reflecting a slow leakage of dithionite into the vesicles or the spontaneous flipping of the NBD-labelled lipids. A linear fit was used to estimate the rate of this process was estimated to be L=(5.4±1.6).10^−5^ s^−1^ (n>160)^32^. For TMEM16K, the leak is >2 orders of magnitude smaller than the rate constant of protein-mediated scrambling and therefore was considered negligible. All conditions were tested side by side with a control preparation of afTMEM16 in standard conditions.

### Statistical analyses

All values are the mean ± s.e.m. Results are expressed as mean ± s.e.m. of *N* (number of experiments). Statistical significance was determined with two-tailed unpaired *t*-test or one-way ANOVA with Bonferroni’s *post hoc* test, as appropriate. For all statistical tests, *P*-values <0.05 were considered significant. SPSS (version 22; SPSS Inc., Chicago, IL, USA) or Excel (2013 Edition, Microsoft, Redmond, WA, USA) were used for statistical analysis.

For scramblase assays, rate constant values reported are the average of 9 individual fluorescence traces from three independent reconstitutions of TMEM16K and afTMEM16.

### Crystallisation and Data Collection

Purified TMEM16K was concentrated to 10-30 mg/ml using a Vivaspin 20 centrifugal concentrator with a 100 kDa molecular weight cut-off. Protein concentration was measured via A280 using a Nanodrop spectrophotometer. TMEM16K was crystallised using both sitting drop vapour diffusion (VD) crystallisation and *in meso* in the lipidic cubic phase (LCP). VD crystals were grown in 0.1M HEPES pH 7.0, 0.1M calcium acetate, 22% (v/v) PEG400, 0.05mM C12E9 at a protein concentration of 10 mg/ml. For LCP crystallisation, 30 mg/ml TMEM16K was combined with 1-(7Z-hexadecenoyl)-rac-glycerol (monoacyl-glycerol 7.9, Avanti Lipids) in a 1:1.5 ratio to form a lipid cubic phase. A 50nl bolus of LCP-reconstituted TMEM16K was dispensed on a glass LCP plate (Marienfeld, Germany) and overlaid with 800 nl of crystallisation solution. TMEM16K crystals grew in 0.1M MES pH 6.0, 0.1M NaCl, 0.1 M CaCl_2_, 30% (v/v) PEG300. For both LCP and VD crystallisation, initial crystals appeared after 1 week and grew to full size within 3-4 weeks. All X-ray diffraction data were collected on the I24 microfocus beamline at the Diamond Light Source (Didcot, UK) from single crystals using a fine phi slicing strategy. Intensities were processed and integrated using XDS^45^ and scaled using AIMLESS^46^.

### Structural Model Building and Refinement

The initial dataset collected on the LCP derived crystals was phased by molecular replacement using PHASER^47^ with the nhTMEM16 structure (PDB: 4WIS) as an initial search model. Phase improvement was performed using *phenix.mr.rosetta*^48^. The final model was built using COOT^49^ and refined using BUSTER^50^ using all data to 3.2Å with appropriate NCS restraints. The final LCP model was subsequently used as a starting model for molecular replacement to solve the structure of detergent-solubilised TMEM16K using an anisotropic dataset collected from a crystal grown using sitting-drop vapour diffusion methods. The anisotropic nature of the VD dataset prevented it being solved using nhTMEM16 (the only high resolution structure at the time) as a molecular replacement search model. Processing of the VD dataset was similar to that for the LCP crystals, with additional anisotropy correction performed using STARANISO^51^. The model geometry of the VD dataset was improved by using LSSR target restraints to the LCP structure during BUSTER refinement in addition to TLS and NCS restraints (Supplementary Table 1).

The final LCP model encompasses residues Ser14 to Gln639. Several loops were poorly order and not modelled; residues 57-67 and part of the α7-α8 loop (residues 472-474) were disordered in chain A. The loops connecting either α5 and α6 or TM3-TM4 were poorly defined in both chains of the dimer and have also not been modelled. The final model also includes three Ca^2+^ ions per monomer along with an additional Ca^2+^ at the N-terminal end of TM10 on the dimer axis. The presence of Ca^2+^ ions at these sites was indicated by peaks in both anomalous difference and PHASER log-likelihood gradient (LLG) maps calculated using a 3.4 Å dataset with high multiplicity collected at a wavelength of 1.65 Å. Two Ca^2+^ ions in each dimer lie at the canonical two Ca^2+^ ion binding site and a third lies at the junction of TM10 and α10. All of these ions have bonds that are less than 2.5 Å to sidechains, suggesting the ions are not hydrated. The fourth Ca^2+^ ion identified in the anomalous difference maps lies on the dimer 2-fold axis, binding to the backbone of the ER loop between TM9 and 10. The > 4 Å interaction distances suggest that this ion is hydrated and is likely to be the result of the high (100 mM) [Ca^2+^] used in the crystallisation conditions (Supplementary Fig. 5h). In addition, elongated density within the dimer interface were interpreted as MAG7.9 lipids.

### Cryo-EM grid preparation

3 μl aliquots of TMEM16K protein purified in UDM at a concentration of 5mg/ml were applied to glow-discharged holey carbon grids (Quantifoil R 1.2/1.3 Cu 300 mesh). Grids were blotted at 80-100% humidity for 3-5 s at 5 °C and plunge-frozen in liquid ethane using a Vitrobot Mark IV, (FEI). Ca^2+^-free TMEM16K grids were prepared in an identical manner using protein solution supplemented with 10 mM EGTA.

### Cryo-EM data collection

Grid optimization and preliminary data set collection was performed using the Tecnai F30 ‘Polara’ transmission electron microscope (Thermo Fisher) at Oxford Particle Imaging Centre (OPIC), Division of Structural Biology, Oxford operated at 300 kV, at liquid nitrogen temperature and equipped with a K2 Summit direct electron detector (Gatan) mounted behind an energy filter (GIF Quantum LS, Gatan). Screening was also performed using a Talos Arctica operating at 200 kV equipped with a Falcon3 detector (Thermo Fisher) at the Central Oxford Structural Microscopy and Imaging Centre (COSMIC) Oxford University. Cryo-EM data collection was carried out on a Titan Krios at COSMIC, operating at 300 kV. Data were acquired with EPU software on a K2 Summit detector (Gatan) mounted behind an energy filter (GIF Quantum LS, Gatan) and operated in zero-loss mode (0-20 eV). Movies (20 frames, 8 s total exposure) were recorded in electron counting mode at a sampling rate of 0.85 Å / pixel and dose rate of 6.6 e-/Å^2^/s (total exposure 52.8 e^−^/Å^2^) with underfocus in the range from 1.0 to 3.0 μm.

### Cryo-EM data processing and model building

All initial processing was carried out in RELION 2.0^52^. Frames in each movie stack were aligned and dose-weighted with MotionCor2^53^. CTF parameters were estimated using CTFFIND 4.0^54^. Dose weighted stacks were subjected to semi-automatic particle picking using Gautomatch (https://www.mrc-lmb.cam.ac.uk/kzhang/Gautomatch/). Particles were picked from a subset of micrographs and 2D classified to produce class averages. The resultant representative class averages were then used as templates in Gautomatch for autopicking the full image set. Particles were subjected to multiple (5-6) rounds of reference-free 2D classification. An initial model was generated *ab initio* in RELION and used as a reference of 3D classification with no symmetry imposed. Particles belonging to the best / highest resolution class(es) were pooled and taken forward into a second round of 3D classification with C2 symmetry. Finally, particles from the highest resolution class were used for auto-refinement. Further static classification with multiple (10) classes did not yield a better resolved reconstruction. The processing schema for the 500 nM Ca^2+^ and Ca^2+^-free datasets are shown in Supplementary Figs. 6 and 7. Local resolution estimation for each final reconstruction was performed with *blocres* in BSOFT55. Post-processing was carried out in RELION using a mask extended by 12 pixels with an additional 12-pixel soft edge that excluded the detergent micelle surrounding the protein.

Model building was carried out manually using the 3.4 Å TMEM16K LCP crystal structure (PDB: 5OC9) as a template. Briefly, chain A of the crystal structure was roughly fitted into the 4.5 Å resolution 500 nM Ca^2+^ post-processed cryo-EM map in UCSF Chimera^56^. Subsequent model building was carried out in COOT^57^. The cytoplasmic domain was rotated into density and then appropriate sub-regions were fit to the density as rigid bodies. TM helices were fitted as rigid units or segmented where appropriate. The C-terminal α10 helix was rotated manually into position and sidechains were fitted using preferred rotamers. The remodelled chain A was superposed onto chain B and globally fitted to the cryo-EM density to create the symmetric dimer. Refinement was carried out at 4.5 Å resolution against the post-processed RELION map (low pass filtered to 4.4 Å and sharpened with a *B*-factor of −260 Å^2^) using *phenix.real_space_refine* using default restraints along with secondary structure restraints. Missing loop regions, not modelled in the crystal structure, were added and the calcium coordination was defined using *phenix.metal-coordination*. The final model encompasses all residues from 14-462 and three Ca^2+^ ions per monomer (Supplementary Table 2).

The 500 nM Ca^2+^ model was used as a starting point for fitting to the Ca^2+^-free reconstruction. The model was initially docked into the map using Chimera and then refined using *phenix.real_space_refine* with global minimisation and morphing using default restraints/constraints and additional secondary structure restraints. A cryo-EM map, low pass filtered to 5.3 Å resolution and sharpened with a *B*-factor of −150 Å^2^, was used for all refinement and model building. The TM1-loop-TM2 region (residues 222-254) required manual rebuilding/fitting in COOT. The calcium ions were also removed from the model. The resultant model was further refined with *phenix.real_space_refine* using global minimisation only (Supplementary Table 2).

### Structural alignments and analysis

Superposition of the LCP and 500 nM Ca^2+^ cryo-EM structures was carried out in PyMol using a subset of residues corresponding to the TM helices that maintain their relative orientation within each dimer (as measured by interhelical angles). Mainchain atoms for residues 192-262 (α6-TM2), 404-452 (TM5-6) and 554-612 (TM9-10) were used for the structural superposition. Overall comparison of the X-ray and cryo-EM structures is further complicated by the fact that there are also small changes at the dimer interface particularly the relative orientation of TM10 and α10 which closely interacts with the adjacent monomer and subtly changes the relative orientation of monomers within each dimer. Consequently for global comparisons we have carried out the superposition across both molecules in the dimer (referred to as the dimer superposition) and for local structural comparisons we have restricted the alignment to the isolated monomers (monomer superposition) using the above residue range. The ‘dimer superposition’ was used to prepare Fig. 5c,d, Supplementary Fig. 8a-f and the Supplementary video. The monomer superposition was used in the preparation of Supplementary Figs. 8g, 8h.

For DYNDOM torsional analysis, which compared the LCP X-ray and cryo-EM monomer structures, a composite set of coordinates was used in which the α10 helix was exchanged between chains (i.e. residues 14-609 from chain A and 611-639 from chain B). Electrostatic potential surfaces were calculated using the PDB2PQR server and PyMol with the APBS_2.1 plugin. Structural figures were generated using PyMol or UCSF Chimera‥

### Molecular dynamics simulations

The atomic coordinates of TMEM16K were converted to their CG representation^58,59^, and built into membranes following the MemProtMD pipeline ^23^. Membranes were built using a mixture of 65% POPE, 32% POPC and 3% POPS.

Atomistic simulations were run following conversion of 100 ns Martini simulation snapshots in a POPC bilayer, using the CG2AT protocol^60^. The Charmm36 forcefield^61^ was used to describe the system. Electrostatics were handled using the Particle-Mesh-Ewald method, and a force-switch modifier was applied to the Van der Waals forces. Dispersion corrections were turned off. Simulations were run for 100 ns using 2 fs time steps with Velocity-rescaling temperature coupling at 325 K using a time constant of 0.1 ps and Parrinello-Rahman semi-isotropic pressure coupling of 1 Bar with a time constant of 2 ps, before extension to 1,100 ns using 4 fs time steps with virtual-sites on the protein and lipids ^62^ Extended simulation were run on the UK HPC facility ARCHER, using time awarded by HECBioSim.

All simulations were run in Gromacs 5.1.2.^63^. Images of proteins were made in PyMOL^64^ or VMD^65^. Graphs were plotted using Python and Matplotlib.

## Supporting information

## Data availability

X-ray crystallography data and structure are deposited in the Protein Data Bank (PDB) with the accession code 5OC9 for the LCP structure. The VD structure data will be deposited. The cryo-EM density map and the atomic coordinates will be deposited in the Electron Microscopy Data Bank (EMDB) and in the Protein Data Bank (PDB).

## Acknowledgements

The SGC is a registered charity (number 1097737) that receives funds from AbbVie, Bayer Pharma AG, Boehringer Ingelheim, the Canada Foundation for Innovation, Genome Canada, Janssen, Lilly Canada, Merck KGaA, Merck & Co., Novartis, the Ontario Ministry of Economic Development and Innovation, Pfizer, São Paulo Research Foundation-FAPESP and Takeda, as well as the Innovative Medicines Initiative Joint Undertaking ULTRA-DD grant 115766 and the Wellcome Trust (106169/Z/14/Z). N.J.G.R. and P.T. were supported by the Wellcome Trust (OXION grant No: WT084655MA) and the British Heart Foundation. R.S. is funded by the British Heart Foundation. P.J.S. and R.A.C. were supported by the Wellcome Trust [208361/Z/17/Z] and the BBSRC [BB/P01948X/1, BB/I019855/1]. T.D.N. was supported by an EPSRC Doctoral Training Centre studentship.

We thank Diamond Light Source Ltd for access to the macromolecular crystallography beamlines and we thank the Diamond Light Source staff for their help with data collection. We acknowledge the Oxford Particle Imaging Centre (OPIC) for providing access to electron microscopes for grid screening. We also acknowledge the Central Oxford Structural Microscopy and Imaging Centre (COSMIC) for providing access to electron microscopes for grid screening and data collection. We acknowledge the use of the UCSF Chimera package56 from the Resource for Biocomputing, Visualisation, and Informatics at the University of California, San Francisco (supported by NIGMS P41-GM103311). Extended molecular dynamics simulations were carried out using computer time on the ARCHER UK National Supercomputing Service (http://www.archer.ac.uk), provided by HECBioSim, the UK High End Computing Consortium for Biomolecular Simulation (hecbiosim.ac.uk), supported by the EPSRC (EP/L000253/1).

We thank all members of the SGC Biotech team, including Claire Strain-Damerell; Kasia Kupinska, Dong Wang and Katie Ellis. We thank all members of the SGC IMP1 group, including Yin Yao Dong and Andrew Quigley. We thank David Eberhardt for help with the electrophysiology experiments. We are grateful to Rod Chalk, Georgina Berridge and Oktawia Borkowska for help with mass spectrometry and Brian Marsden and David Damerell, James Bray, James Crowe and Chris Sluman for bioinformatics support. We thank Frank von Delft, Tobias Krojer and Beth Maclean for assistance with crystallography infrastructure. We thank Owen Vickery for implementation of the lipid virtual sites and Mark Sansom for discussions. We thank Errin Johnson and Adam Costin (COSMIC) for assistance with grid screening and data collection. We thank Andrea Nemeth for critical reading of the manuscript. Molecular graphics were performed with UCSF Chimera, developed by the Resource for Biocomputing, Visualization, and Informatics at the University of California, San Francisco, with support from NIH P41-GM103311.

## Author Contributions

S.R.B., A.C.W.P. and M.E.F. contributed equally to this project. S.R.B., A.C.W.P., R.S., P.T., P.J.S., A.A. and E.P.C. designed the project. S.R.B. optimised and prepared protein samples, obtained crystals and prepared cryo-EM grids with C.A.S., assisted by A.T. and A.C‥ Q.W. assisted with lipid binding studies. S.R.B., A.C.W.P. and E.P.C. screened crystals and collected X-ray datasets. S.R.B., C.A.S., J.T.H., A.C.W.P. and E.P.C. screened grids and collected cryo-EM data. S.R.B. and A.C.W.P. determined the structures. L.S. and S.M.M.M. performed pilot studies for protein production and produced baculovirus infected cells. Work by L.S., and S.M.M.M. was supervised by N.A.B.-B‥ J.D.L. was involved in pilot studies for protein production. N.J.G.R., C.M.T. and P.T. performed and analysed patch-clamp experiments and N.J.G.R., supervised by P.T., performed the immunocytochemistry. M.E.F. performed the lipid scramblase and bulk ion channel experiments, supervised by A.A. S.R.B attempted single channel e-physiology experiments (not shown as the results were inconclusive), supervised by R.S. Molecular dynamics simulations were designed, performed and analysed by R.A.C., T.D.N. and P.J.S‥ E.P.C. was responsible for the overall project design, data collection, supervision and management. S.R.B., A.C.W.P., M.E.F., N.J.G.R., T.R.N., R.A.C., R.J.S., J.T.H., P.T., A.A and E.P.C. analysed the data and prepared the manuscript.

