## Supplementary material for "The structural basis of lipid scrambling and inactivation in the endoplasmic reticulum scramblase TMEM16K"

### Supplementary Figures

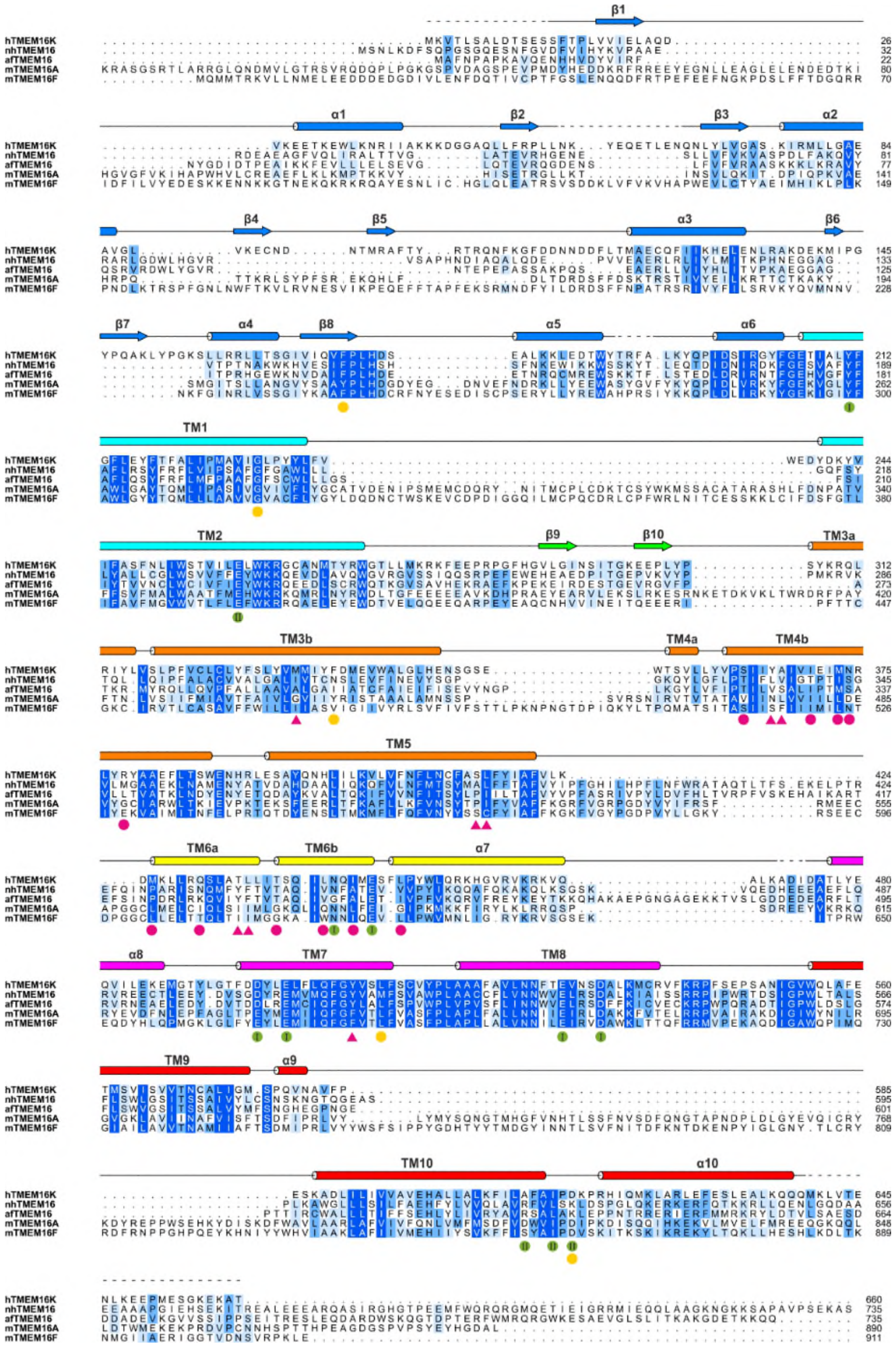

**Supplementary Fig. 1** Structure-based sequence alignment of human TMEM16K (UniProt: Q9NW15, ANO10_HUMAN), mouse TMEM16A (Q8BHY3, ANO1_MOUSE), mouse TMEM16F (Q6P9J9, ANO6_MOUSE) and TMEM16 homologs from fungal species *Nectria haematococca* and *Aspergillus fumigatus*. Conserved and partially conserved residues are highlighted in dark blue and light blue respectively. Residues involved in TM6-8 $Ca^{2+}$-binding site and TM10/α10 $Ca^{2+}$-binding site are marked with green dots. Known disease mutations are marked with orange dots. Residues located in the scramblase groove that were identified as forming contacts with lipids in MD simulations are shown as magenta circles and residues in the neck region are shown as magenta triangles. Secondary structure annotation is derived from the LCP TMEM16K crystal structure and are coloured as described in Fig. 2b.

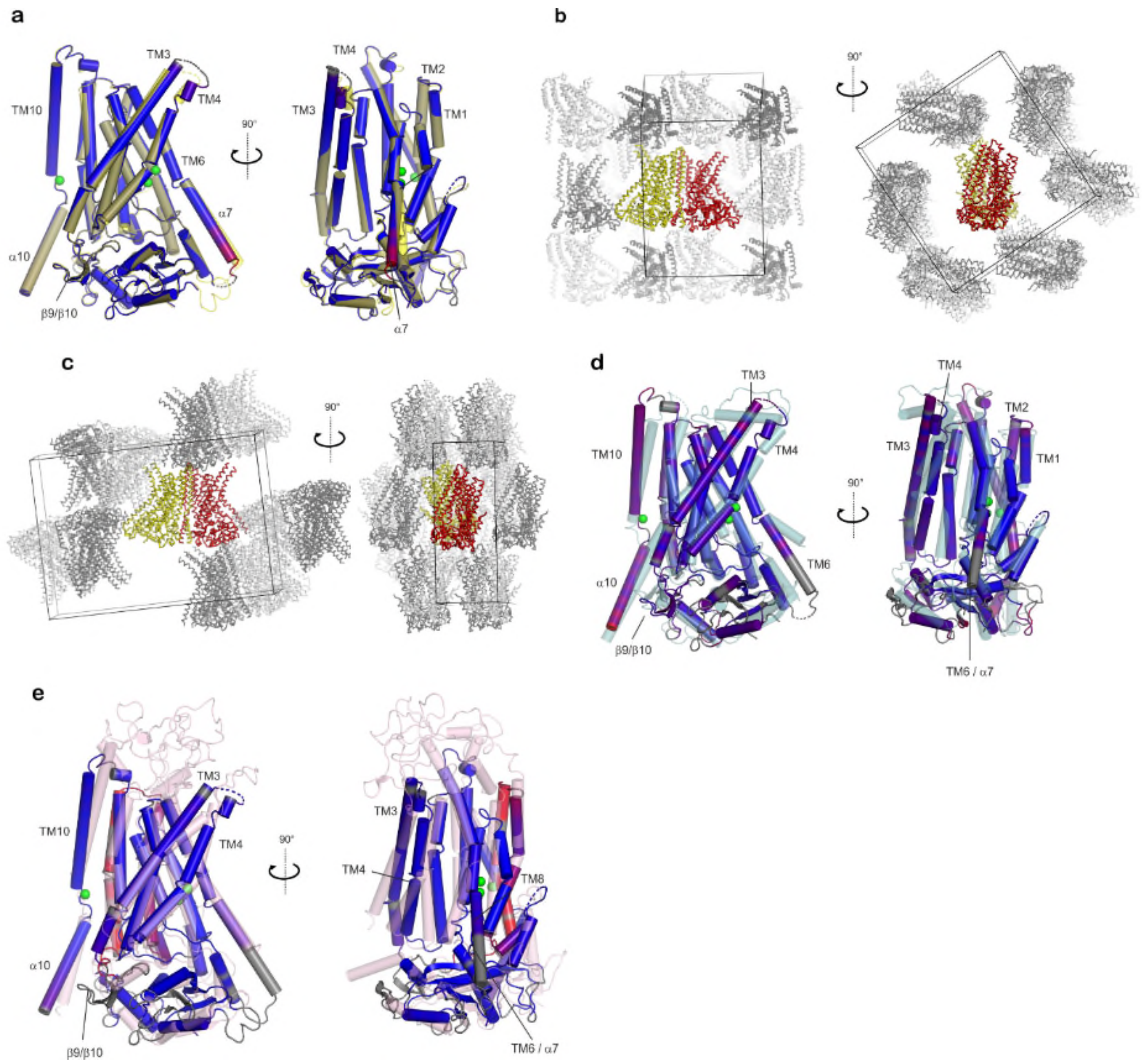


**Supplementary Fig. 2** Superpositions of TMEM16K with various homologs and crystal packing. **a** Superposition of TMEM16K crystal structures. The monomer from the vapour diffusion derived crystal structure is coloured as semi-transparent gold. The LCP monomer is coloured by the root mean square differences between the structures, with blue representing those most closely aligned and red the most distant. Bound calcium atoms are depicted as green spheres. **b-c.** Crystal packing within (**b**) vapour diffusion and **c** LCP crystal forms. TMEM16K is depicted as yellow (chain A) and red (chain B) Cα traces. The unit cell is depicted as a black cube, with symmetry mates depicted as light and dark grey Cα traces. For clarity, some symmetry mates which obscure view of the central molecule are not shown, though all crystal contacts are represented in the figure. **d-e** Superposition of TMEM16K (coloured by RMS deviation as described in Supplementary Fig. **2a**, with (**d**) nhTMEM16K, coloured white, or (**e**) mTMEM16A, coloured gold. Bound calcium in the TMEM16K structures depicted as green spheres.

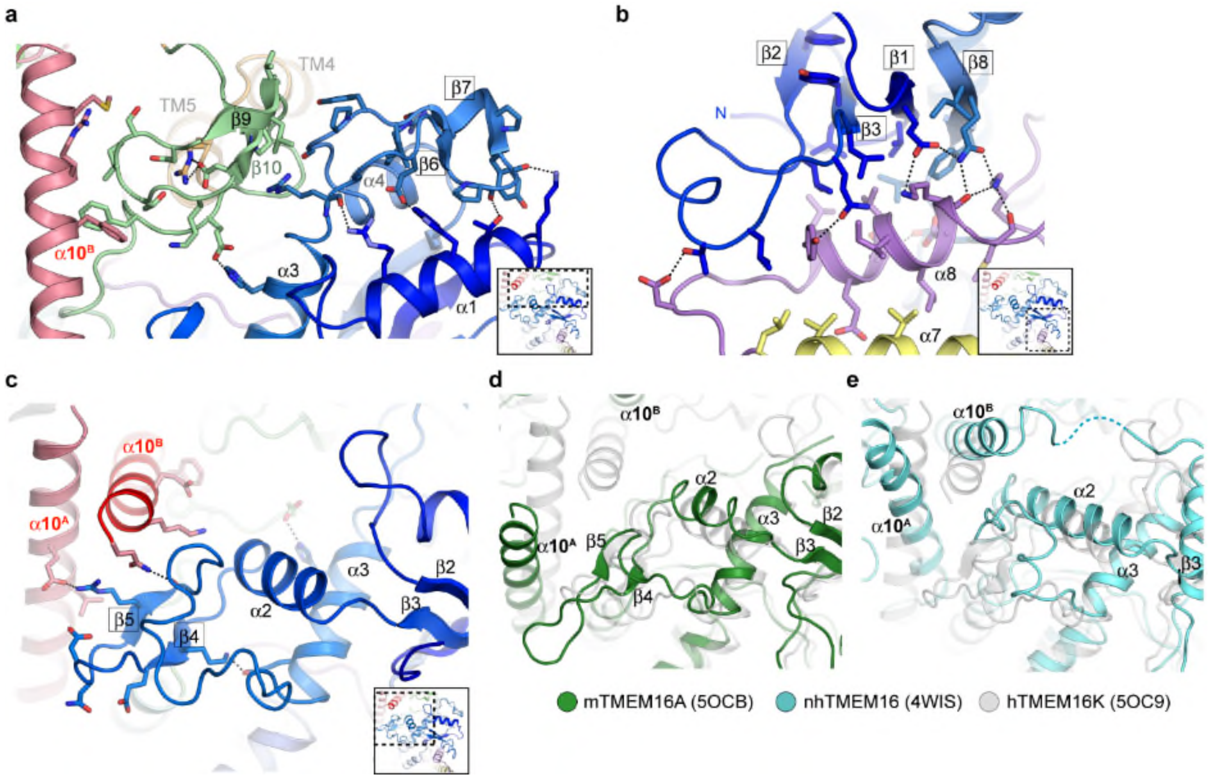


**Supplementary Fig. 3** Key interactions between the TMEM16K N-terminal cytoplasmic domain (NCD) the TM domain extensions and α10. **a** β6-β7 hairpin and the associated loops interactions with the β9-β10 hairpin on the TM domain. **b** four stranded β-sheet (β1-β3, β8) is tightly backed against α8, part of an α-helical hairpin formed by α7 and α8, which are extensions of TM6 and TM7. **c** Between β3 and α3 there is a region where the TMEM16K, mTMEM16A and nhTMEM16 structures differs significantly, TMEM16K having a relatively short α3 and a β4-β5 hairpin that interacts directly with the C-terminal $\alpha10^{A}$ and $\alpha10^{B}$ helices. **d-e** Corresponding region in (**d**) mTMEM16A (green) and (**e**) nhTMEM16 (cyan).

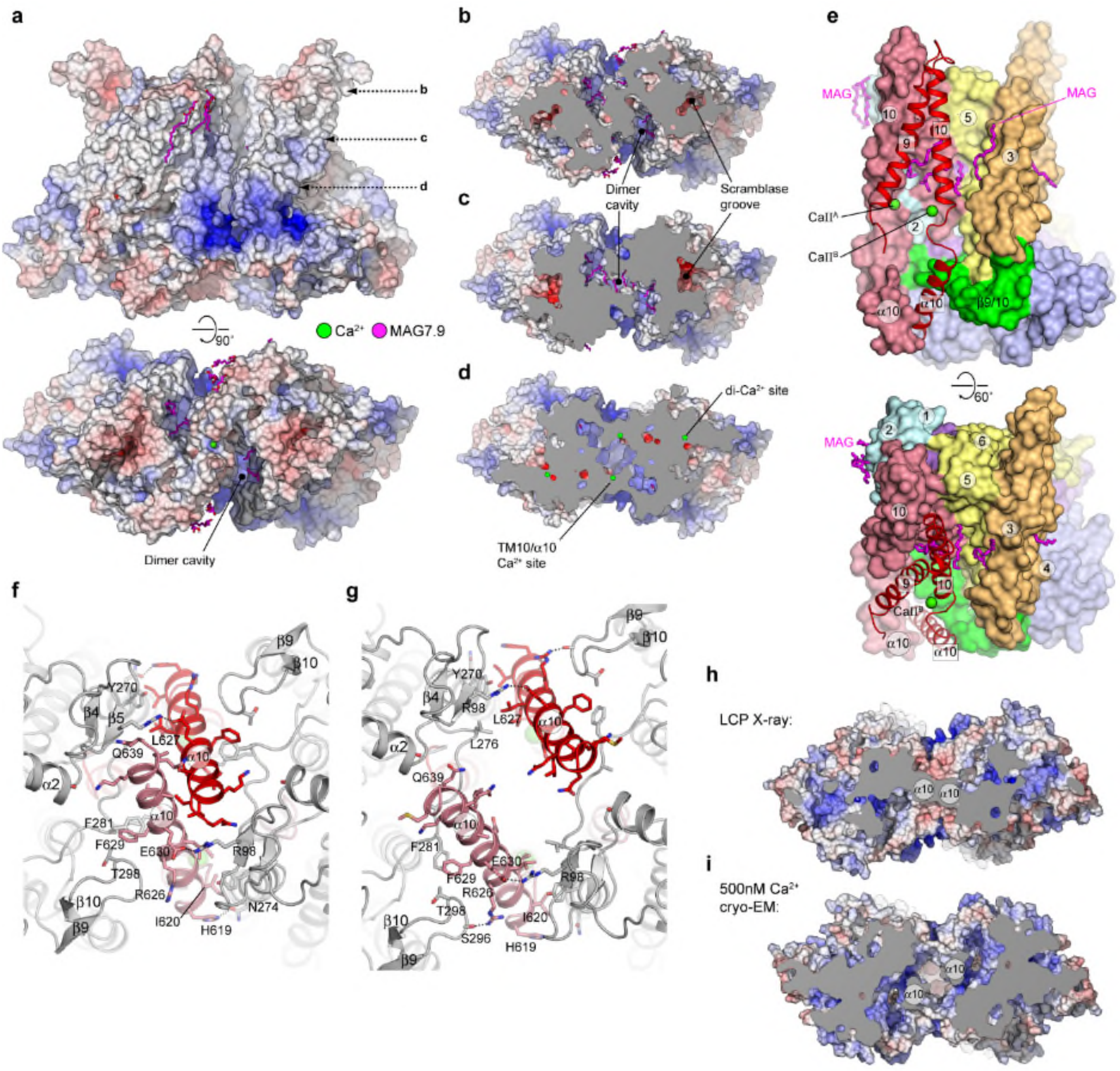


**Supplementary Fig. 4** The TMEM16K dimer interface and the TM10-α10 Ca $^{2+}$-binding site. **a** Molecular surface for TMEM16K LCP X-ray structure coloured by electrostatic potential (+/- 10kT/e). **b-d** Sliced molecular surface viewed from ER luminal face showing dimer interface cavity. Approximate locations of sliced view indicated in **a.** **e** Cut-through of TMEM16K's dimer interface. Chain A is represented as a space-filled model with TM9-10 and α10 from Chain B depicted as ribbon. The TM10/α10 calcium binding sites $CaII^A$ and $CaII^B$ are derived from Chains A and B respectively and calcium ions are depicted as green spheres. MAG lipids are coloured magenta. **f-g** Comparison of the interactions of α10 region of dimer interface for the (**f**) LCP X-ray and (**g**) 500 nM $Ca^{2+}$ cryo-EM structures. The dimer interface is viewed from the cytoplasm. **h-i** Sliced molecular surfaces for (**h)** LCP X-ray and (**i**) 500 nM $Ca^{2+}$ cryo-EM structures highlighting considerable reduction in the closed scramblase's dimer interface due to repositioning of α10 helices. Molecules are viewed from the cytoplasmic face.

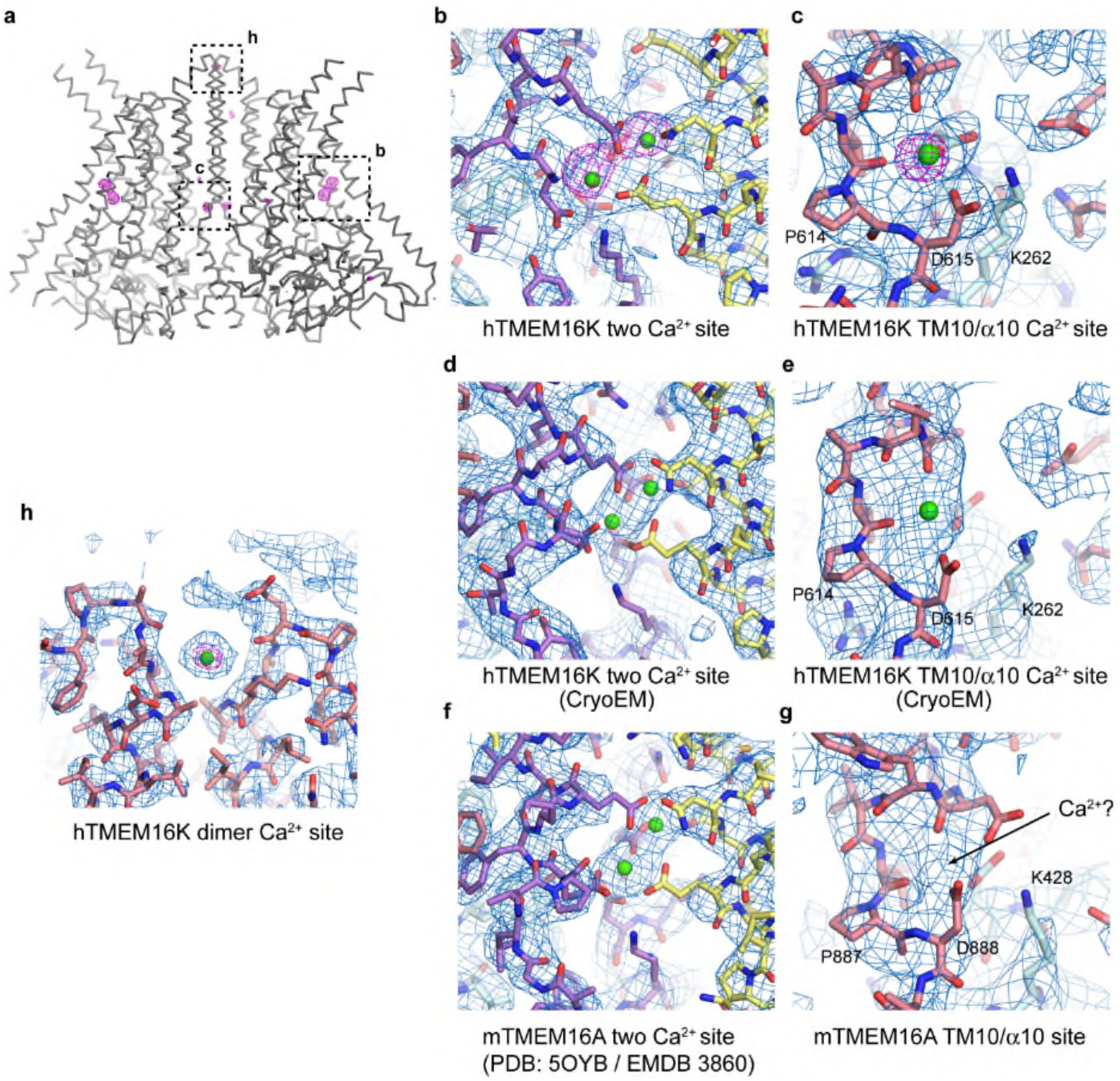


**Supplementary Fig. 5** Electron density maps at the $Ca^{2+}$ binding sites in TMEM16K and TMEM16A. **a** PHASER single anomalous diffraction log-likelihood gradient (LLG) map (magenta mesh, contoured at 4.5 σ) calculated from data collected at a wavelength of 1.65Å and overlaid on a Cα trace of the final LCP crystal structure. Clear peaks are observed for all assigned $Ca^{2+}$ atoms. **b-c,** Detailed view of **b** two $Ca^{2+}$ site **c** TM10-α10 $Ca^{2+}$ binding site from the LCP crystal structure. Final BUSTER 2FoFc electron density (blue mesh) contoured at 1σ and PHASER LLG map (magenta mesh, contoured at 5.5σ) overlaid on final refined model. **d-e** Corresponding density in the 4.6 Å 500 nM $Ca^{2+}$ cryo-EM map around **d** the TM6-8 two $Ca^{2+}$ and **e** TM10 / α10 $Ca^{2+}$ site. The map is contoured at 3σ. **f-g** Cryo-EM maps around **f** the two $Ca^{2+}$ site and **g** the TM10-α10 region for the 3.75Å mouse TMEM16A structure (PDB: 5OYB / EMDB 3860). The map is contoured at 7.5σ. **h** Third $Ca^{2+}$ site located on the dimer axis between TM10s in the LCP crystal structure. LLG density map (magenta mesh) is contoured at 4σ.

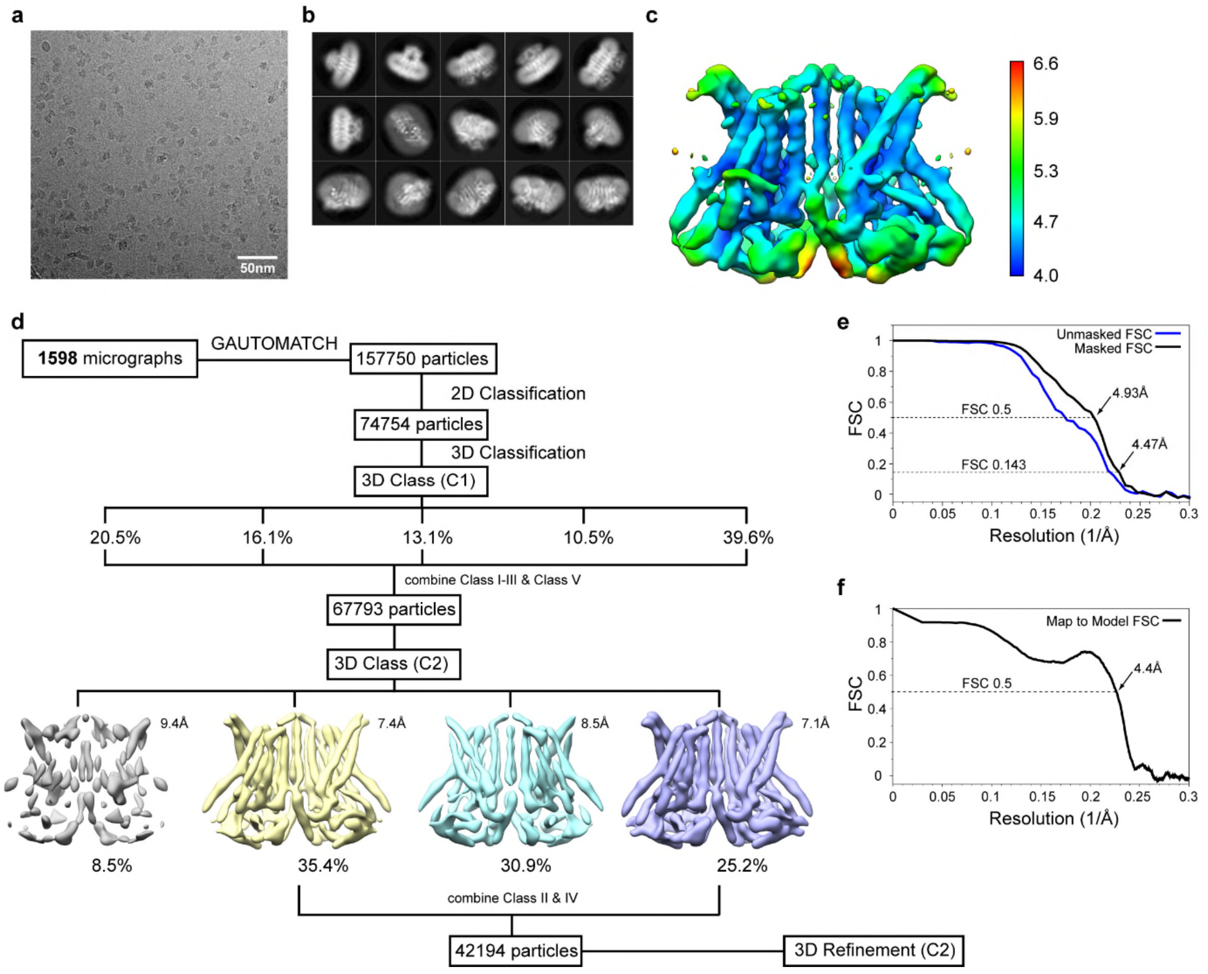


**Supplementary Fig. 6** Cryo-EM details of $Ca^{2+}$-bound TMEM16K. **a** Representative micrograph. **b** Representative 2D class averages (200px box). **c** Local resolution map calculated using *blocres*. **d** Workflow for data processing in RELION. **e** FSC analysis for final 3D reconstruction. **f** Map to model FSC for refined model as calculated by *phenix.mtriage.*

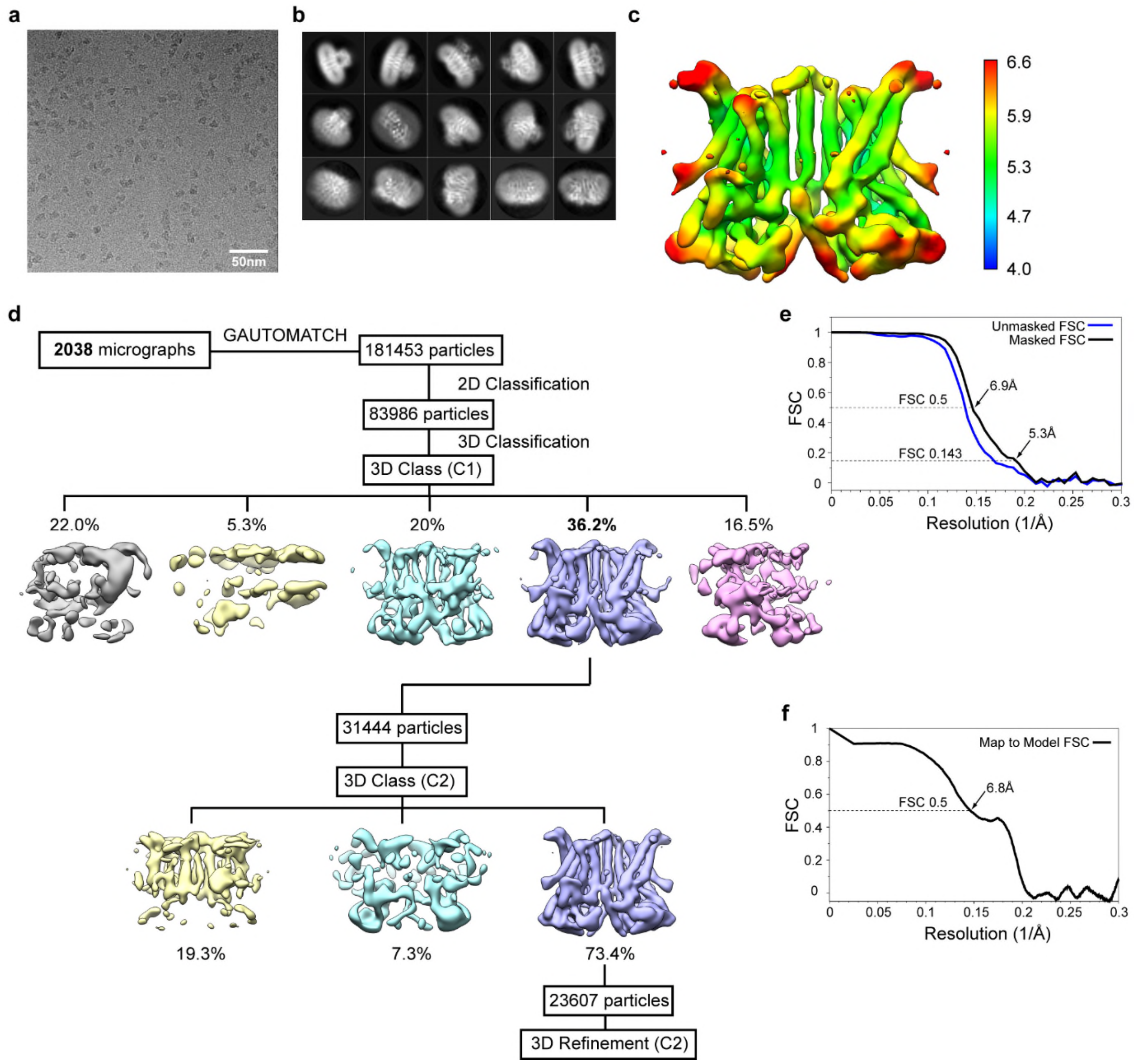


**Supplementary Fig. 7** Cryo-EM details for $Ca^{2+}$-free / EGTA TMEM16K. **a** Representative micrograph. **b** Representative 2D class averages (200px box). **c** Local resolution map calculated using *blocres*. Resolution colour scale is the same as Extended Data Fig. 6c. **d** Workflow for data processing in RELION. **e** FSC analysis for final 3D reconstruction. **f** Map to model FSC for refined model as calculated by *phenix.mtriage.*

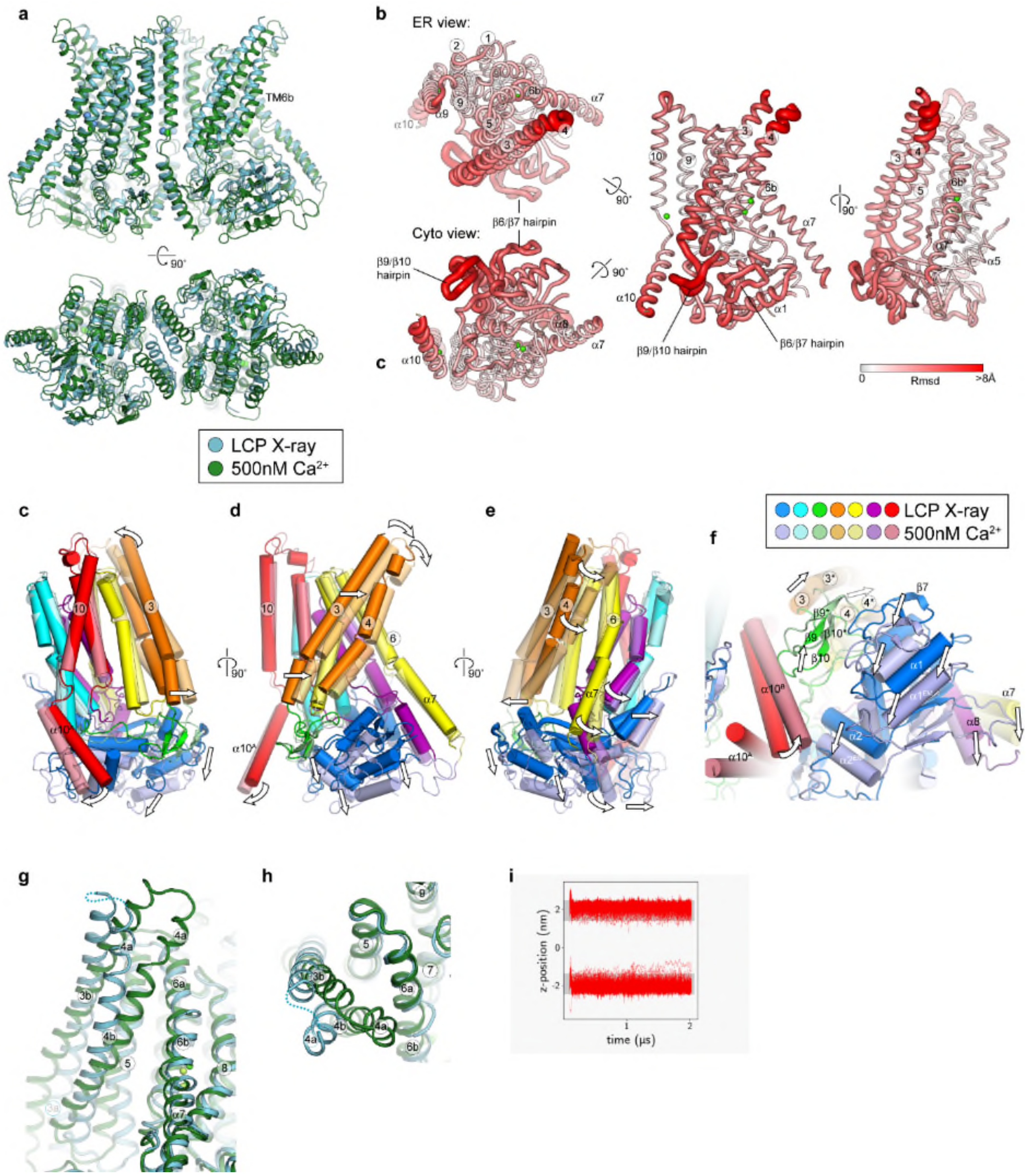


**Supplementary Fig. 8** Conformational changes between the crystal and cryo-EM structures. **a** Overall comparison of the LCP X-ray (blue) and 500nM $Ca^{2+}$ cryo-EM structure (green). **b** Global conformational differences between LCP X-ray and 500nM $Ca^{2+}$ cryo-EM monomer structures. Schematic representations of the rms deviation (rmsd) in mainchain atomic positions mapped onto the monomer structure. Monomers from each structure were superposed using all atoms with LSQKAB (CCP4). The monomer is viewed from either the luminal or cytoplasmic face (*left*), perpendicular to the membrane normal (*middle*) and onto scramblase groove face (*right*). The thickness and colour of the tube reflects the magnitude of the rmsd between the two structures. **c-f**, Conformational differences between LCP X-ray (5OC9) and 500nM $Ca^{2+}$ structures. Schematic representation looking onto (**c**) dimer face, (**d**) perpendicular to the membrane normal, (**e**) the scramblase groove and (**f**) cytoplasmic domain. The structures were superposed in the context of the full dimer using the scaffold region alone (see Methods). The X-ray structure is coloured in darker shades and the 500 nM $Ca^{2+}$ cryo-EM structure in lighter shades. **g-h**, Conformational differences in the vicinity of the scramblase groove between TM3-TM7 viewed (**g**) perpendicular to the groove (**h**) looking down the groove from the luminal face. **i** MD simulations, showing lack of movement of lipids across the membrane with the closed groove.

### Supplementary Tables

**Supplementary Table 1** X-ray crystallography data, refinement and model statistics.

| | TMEM16K LCP | TMEM16K VD |
|---|---|---|
| **PDB Code** | 5OC9 | |
| **Data Collection** | | |
| Beamline | I24 | I24 |
| Space group | $P2_12_12_1$ | $P2_12_12_1$ |
| Crystallisation conditions | 0.1M MES pH 6.0, 0.1M NaCl, 0.1M $CaCl_2$, 30% PEG300 | 0.1M HEPES pH 7.0, 0.1M Ca Acetate, 22% PEG 400, 0.05mM C12E9 |
| Cell dimensions | | |
| *a*, *b*, *c* (Å) | 59.15, 153.59, 218.78 | 113.16, 152.27, 153.70 |
| α, β, γ (°) | 90, 90, 90 | 90, 90, 90 |
| Resolution [Å][1] | 3.2 (3.20-3.28)[1] | 3.5 (3.59-3.50)[1] |
| Resolution limits [Å][2,3] | 4.00, 3.43, 3.20 | 3.66, 5.86, 3.63 |
| | (3.68, 3.25, 3.20)[3] | (3.50, 5.16, 3.50)[3] |
| Nominal Resolution [Å][3] | 3.39 | 3.97 |
| $CC_{1/2}$ | 0.990 (0.535) | 1.00 (0.352) |
| $R_{meas}$ | 0.318 (1.398)[1] | 0.201 (2.200)[1] |
| $R_{pim}$ | 0.139 (0.610) | 0.103 (1.104) |
| $I / \sigma I$ | 5.9 (1.4)[1] | 5.6 (0.8)[1] |
| Completeness [%] | 99.9 (99.9)[1] | 99.6 (99.8)[1] |
| Redundancy | 5.1 (5.2)[1] | 3.7 (3.8)[1] |
| **Refinement** | | |
| Resolution (Å) | 76.79 – 3.2 | 46.28 – 3.65 |
| No. reflections (free) | 33824 (1699) | 29961 (1492) |
| $R_{work}$ / $R_{free}$ | 22.9 / 24.5 | 25.74 / 27.0 |
| No. atoms | | |
| Protein | 9806 | 9630 |
| Other | 332 | 6 |
| *B*-factors (Å$^2$) | | |
| Protein | 57.76 | 137.15 |
| Other | 54.04 | 122.32 |
| R.m.s. deviations | | |
| Bond lengths (Å) | 0.008 | 0.008 |
| Bond angles (°) | 0.87 | 0.90 |

[1] Values in parentheses are statistics for highest resolution shell

[2] Anisotropic resolution limits along each of the three principal directions as defined by AIMLESS based on Mn (I/sd(I)) > 2

[3] Values in parentheses are resolution limits in each direction based on half dataset correlation > 0.5 ($CC_{1/2}$).

[4] Nominal resolution is defined based on overall Mn (I/sd(I)) > 2 as estimated by AIMLESS.

**Supplementary Table 2** Cryo-EM data, refinement and model statistics.

| **Cryo-EM data** | 500 nM $Ca^{2+}$ | $Ca^{2+}$-free |
|---|---|---|
| **Data Collection:** | | |
| Voltage (kV) | 300 | |
| Defocus range (μm) | -1.1 to -3.0 | |
| Pixel size (Å) | 0.85 | |
| Electron dose ($e^-/Å^2$) | 52.8 | |
| Dose rate ($e^-/Å^2/s$) | 6.6 | |
| Number of micrographs (used) | 1975 (1577) | 2083 (2038) |
| Particles (initial)[1] | 157750 | 143308 |
| Particles (final, %) | 42194 (27) | 23607 (16) |
| **Reconstruction:** | | |
| Number of particles | 42194 | 23607 |
| Symmetry | C2 | C2 |
| Resolution (unmasked, Å)[2] | 4.59 | 6.3 |
| Resolution (masked, Å)[2] | **4.47** | **5.31** |
| Map sharpening *B*-factor ($Å^2$) | -260 | -383 |
| **Model composition:** | | |
| Non-hydrogen atoms | 10044 | 9808 |
| Protein residues | 1264 | 1234 |
| **Refinement:** | | |
| Resolution (Å) | 4.5 | 5.3 |
| Map sharpening factor ($Å^2$) | -260 | -150 |
| Fourier Shell Correlation (FSC)[3] | 0.7943 | 0.8207 |
| d_FSC (Map vs. Model)[4] | 4.18 (4.41) | 5.09 (6.82) |
| **Rms deviations:** | | |
| Bonds (Å) | 0.006 | 0.005 |
| Angles (°) | 0.854 | 0.792 |
| **Molprobity validation:** | | |
| Clashscore, all atoms | 2.25 | 3.18 |
| Molprobity score | | |
| Ramachandran Plot (% favoured) | 96.01 | 95.11 |
| Ramachandran Plot (% allowed) | 3.99 | 4.89 |
| Ramachandran Plot (% outliers) | 0.0 | 0.0 |

[1] Particles after one cycle of 2D classification to remove non-particles
[2] Based on FSC 0.143 threshold
[3] CC_mask from *phenix_real_space_refine*
*4 d_fsc resolution estimate FSC=0.143 (FSC=0.5) from phenix.mtriage*
